# Graph Structured Neural Networks for Perturbation Biology

**DOI:** 10.1101/2024.02.28.582164

**Authors:** Nathaniel J. Evans, Gordon B. Mills, Guanming Wu, Xubo Song, Shannon McWeeney

## Abstract

1

Computational modeling of perturbation biology identifies relationships between molecular elements and cellular response, and an accurate understanding of these systems will support the full realization of precision medicine. Traditional deep learning, while often accurate in predicting response, is unlikely to capture the true sequence of involved molecular interactions. Our work is motivated by two assumptions: 1) Methods that encourage mechanistic prediction logic are likely to be more trustworthy, and 2) problem-specific algorithms are likely to outperform generic algorithms. We present an alternative to Graph Neural Networks (GNNs) termed *Graph Structured Neural Networks* (GSNN), which uses cell signaling knowledge, encoded as a graph data structure, to add inductive biases to deep learning. We apply our method to perturbation biology using the LINCS L1000 dataset and literature-curated molecular interactions. We demonstrate that GSNNs outperform baseline algorithms in several prediction tasks, including 1) perturbed expression, 2) cell viability of drug combinations, and 3) disease-specific drug prioritization. We also present a method called *GSNNExplainer* to explain GSNN predictions in a biologically interpretable form. This work has broad application in basic biological research and pre-clincal drug repurposing. Further refinement of these methods may produce trustworthy models of drug response suitable for use as clinical decision aids.

**Availability and implementation:** Our implementation of the GSNN method is available at https://github.com/nathanieljevans/GSNN. All data used in this work is publicly available.

## 2 Introduction

The development of new therapies is a complex and multifaceted task that requires a detailed understanding of disease and the effect of a potential treatment. To develop cancer therapies, for instance, it is critical to know the state of a tumor (e.g., cancer driving mechanisms) as well as the impact a therapy will have on malignant cells, the tumor microenvironment and the host ecosystem. The precision medicine paradigm adds additional layers of complexity due to the need to tailor treatment to disease characteristics. Computational modeling is ubiquitous in this research and can be used to accelerate the discovery of new drug candidates, to understand complex disease mechanisms, and for use as clinical decision aids. Consequently, the accuracy and usefulness of modeling strategies is crucial for effective research.

The field of cancer drug response has developed numerous models for the prediction of simple phenotypic readouts such as cell viability, and these models have been used widely for drug prioritization and basic research. Cell viability, however, can be mediated by a multitude of biological processes and can vary significantly depending on the celluar context or disease. Due to these aspects, cell viability is unlikely to provide the information necessary to characterize the underlying mechanism of response. The systems biology perspective, on the other hand, focuses on modeling the complex and high-dimensional behavior of a disease or process and can be used to understand both outcome and mechanism. Accurate computational models of systems biology have broad application in research and translational medicine.

Perturbation biology is a discipline within systems biology that studies how minor alterations, such as the introduction of a ligand, drug, or genetic lesion, can lead to significant functional changes in an organism [43, 64, 51]. This field investigates the broad effects of such perturbations, often by comparing molecular features before and after a change in a biological system. The emergence of high-throughput sequencing technologies has been pivotal in this area, enabling the extensive measurement of various ‘omics’ levels, such as proteomics, transcriptomics, and genomics. Key technologies in this domain include RNA sequencing [48], the L1000 assay [85], and reverse-phase protein array (RPPA) [19]. These tools are crucial for exploring the systemic response of an organism to perturbations, providing a comprehensive view of the intricate network of biological interactions and the response to a perturbation. The molecular changes caused by a perturbation may be post-translational (e.g., phosphorylation or ubiquitation of a protein), epigenetic (e.g., methylation changes or chromatin remodeling) or expression changes (RNA or protein). System responses begin in an unperturbed state and develop over time. Typically, measurement assays capture these responses at a single time-point, but to fully characterize temporal dynamics, multiple assays are needed, often at a prohibitive cost. This limitation can be addressed by employing computational models, which extrapolate the full temporal response from limited data. However, the critical factor in this approach is the precise timing of the assay measurement. Tailoring measurement times to specific biological processes can be challenging due to their varied rates [81].

In our research, we adopt a premise of *drug-target perturbation response*. Within this premise, a drug molecule binds to one or more proteins and causes a conformational change that alters the protein function and leads to a cascade of physically interacting molecular elements (proteins, RNA, ligands, etc.). These signaling cascades often culminate in the activation of transcription factors (TFs). TFs regulate gene expression by binding to sequence-specific DNA genes and increasing or inhibiting transcription. Activation of a TF can also initiate more complex transcriptional programs via gene regulatory networks (GRN), which comprise a web of interacting transcriptional regulators and may include microRNAs (miRNA), long non-coding RNA (lncRNA) and transcription factors. Additionally, GRNs do not necessarily require regulator-regulator interactions and may have intermediary post-translational signaling (e.g.,regulator→protein-protein signaling→regulator). In summary, our *drug-target* premise envisions a complex network of physically interacting molecular entities that characterize the systemic response to a perturbation.

Drug-protein, protein-protein and gene-regulatory interactions have been studied, categorized, and made available through many public knowledge bases including STRING [88], Reactome [28], Omnipath [90], Targetome [12] and STITCH [47]. Cellular context, which may depend on cell type, disease, species, or genetic background, will define the subset of active interactions in a given biological system. Most database resources do not specify in which cellular context an interaction is active, and therefore reported molecular interactions should be considered as a set of *possible* interactions across all cellular contexts. The unique activity of molecular interactions within a given context, sometimes called the “edgotype” [75], can be attributed to many mechanisms. The binding affinity between a drug and a protein can vary due to gene mutations [30] and the functional effect of a drug will depend on the concentration of the target protein. Differences in protein expression between cellular contexts can mediate different responses to a drug. Precision oncology often takes advantage of these differences by developing drugs that preferentially target proteins that are overexpressed in certain cancers, such as the use of HER2 targeting tyrosine kinase inhibitors (TKI) in HER2+ breast cancer [79]. The contextual “edgotype” can also be mediated by the expression of key molecular entities, and gene mutations may result in nonfunctional protein products. Some gene mutations can prevent or encourage specific protein-protein interactions (PPI) [75]. Differences in expression and genetic background affect cell signaling and can lead to divergent contextual responses to the same drug. The ability of a TF to regulate downstream genes (referred to as the “regulon” of a TF) depends on the expression and state of the TF and other co-regulatory proteins. Additionally, chromatin organization, methylation, and gene mutations can affect the ability of a TF to bind to its DNA targets [34, 52, 75]. Although there has been considerable research characterizing molecular interactions, many knowledge bases (including those used in this study) are not complete, and many important interactions may be missing. Drug-target interactions, for example, are notoriously sparse and many bioactive compounds do not have a known protein target: An estimated 7-18% of FDA-approved drugs do not have known molecular targets [63].

Machine learning (ML) methods are often evaluated by their ability to accurately predict the outcome variable. Even accurate models, however, may be insufficient for many use-cases if they are not trustworthy. The National Institute of Standards and Technology (NIST) defines the characteristics of *Trustworthy* Artificial Intelligence (AI) as [2]:

- Validity and Reliability
- Safety
- Security and Resilience
- Accountability and Transparency
- Explainability and Interpretability
- Privacy
- Fairness with Mitigation of Harmful Bias

These characteristics are intended to provide guidelines for the development of artificial intelligence models that can be trusted to perform in a beneficial manner. Complex and powerful modeling strategies, such as deep learning, often lack many of these aspects of trustworthiness. In particular, most deep learning algorithms are considered a “black box,” which refers to the inability of humans to explain or interpret predictions. Not knowing how a prediction is made can lead to poor generalization or unexpected behavior in new settings, and making decisions based on “black box” model predictions may result in harm.

The development of reliable and high-performance perturbation response algorithms will pave the way for highly impactful applications in translational and research settings. Pre-clinical precision oncology research can use models of perturbation biology to prioritize therapeutic drug candidates for further study. Basic science can use interpretable and explainable models to understand the nuanced and complex behavior of biological systems. Future clinicians may be able to use trustworthy models of drug response to choose optimal patient treatment options based on tumor or patient characteristics.

### 2.1 Ideal Modeling Requirements of Perturbation Biology

In this section, we highlight several (but not necessarily all) important aspects of signaling that an effective modeling strategy should be able to capture.

#### Molecular state

Many molecular entities, such as proteins, RNA, and DNA, have a contextual *state*, which describes its unique behavior and interactions within the system; the contextual state is likely to vary by disease, patient, and cell type. For example, a protein’s state may include aspects such as expression, post-translational modifications, mutational status, or complex co-factors. An ideal algorithm should be able to infer molecular entity state from contextual features (e.g., ‘omics) and appropriately mediate the prediction logic to align with true contextual signaling patterns.

#### Source awareness

The signaling behavior of molecular entities may vary based on the source of upstream signaling. For example, G protein-coupled receptors (GPCR) and receptor tyrosine kinases (RTK) are known to exhibit *ligand bias*, which describes unique signaling patterns dependent on the specific ligand or drug that binds to the receptor [82, 44]. The behavior of a protein can also depend on the source of the upstream signal and may have stimulatory or inhibitory regulators that cause distinct downstream behavior. An ideal algorithm should be able to delineate specific input signals to a molecular entity and simulate different downstream signaling based on the source.

#### Signal latency

Perturbation response evolves over time, and molecular relationships have different rates at which they progress. For example, we expect most post-translation modifications to be much faster than transcription and translation [81]. The rate of PPI signaling is characterized by the association of proteins and is mediated by molecular diffusion rates and conformational kinetics [80]. The rates of specific molecular interactions are, to our knowledge, not available in any resource and therefore an ideal algorithm should be able to infer signal latencies from the training data.

#### Nonlinear multivariate input-output relationships

Many molecular interactions in cellular signaling are known to exhibit *amplification*, where a small input signal may lead to a large output signal, and are likely to be non-linear based on computational models [35, 83]. Additionally, protein-protein signaling often exhibits unique behavior dependent on multiple upstream signals and can be modeled as logic gates. For example, proteins may require two or more upstream signals (AND gate) to be activated, can be activated by multiple signals (OR gate), or a protein may need the absence of an inhibitory signal (NOT gate). Temporal attenuation due to feedback loops or autoregulation can lead to nonlinear temporal relationships in signaling. These characteristics suggest that an ideal model should be able to learn multivariate and nonlinear relationships between input and output signals.

### 2.2 Current limitations of traditional Neural Networks applied to cellular signaling

An artificial neural network (NN) [76, 77] is a machine learning algorithm that has been applied to many domains with great success. NNs have been shown to be *universal approximators*, meaning that they can learn any continuous function if given sufficient data and resources [38]. In most applications, however, machine learning has limited training data and NNs suffer from *overfitting*, which is the tendency of models to accurately predict within training data but generalize to new data poorly. There are many ways to address overfitting, such as curating larger datasets, performing data augmentation [65] or using regularization methods such as weight decay or dropout [84]. Despite extensive research in this area, overfitting remains a challenge in many prediction tasks. Processes that involve high-dimensional inputs or outputs, or characterize complex systems, are likely to have an immense *hypothesis space*^1^ and finding the appropriate hypothesis ^2^ can be exceptionally challenging, especially in limited or noisy data settings.

The *no free lunch theorem* (NFL) states that the performance of all optimization algorithms, when averaged over all possible problems, will be equivalent and suggests that developing unique optimization algorithms tailored to specific problems is key to improving performance [97]. A strategy to create algorithms tailored to specific problems involves integrating inductive biases. This approach enables the model to favor certain solutions over others, regardless of the data presented. [8]. For example, when working with image data, a rational assumption is that nearby pixels are more relevant to each other, and convolutional neural networks (CNN) [3] incorporate this inductive bias by applying a shared function^3^ to local regions of the image, which can reduce the number of trainable parameters and improve performance. Similarly, in natural language processing (NLP), the Transformer model [92] is designed to take advantage of a shared dictionary of tokens and sequence order. Both CNNs and Transformers have been shown to significantly outperform traditional neural networks in their respective problems [92, 3]. Researchers can use these domain-specific characteristics to incorporate relevant assumptions about the data into learning algorithms, which markedly narrows the hypothesis space, often leading to improved performance and usefulness.

The scientific community has put great effort into developing well-supported theories of drug response and perturbation biology. Including these assumptions into a perturbation biology learning algorithm is likely to lead to far more mechanistic, accurate, and useful predictive models; unfortunately, traditional neural networks do not have a convenient means to incorporate this prior knowledge.

### 2.3 Current limitations of Graph Neural Networks applied to cellular signaling

Graph Neural Networks (GNN) are designed to learn the relationships between graph structures and an outcome variable. For instance, the CORA citation network characterizes academic manuscripts (nodes) that cite each other (edges) and GNNs can be used to accurately infer manuscript research domains [45]. Foundational GNNs include the Graph Convolutional Network (GCN) [45], the Graph Isomorphism Network (GIN) [98] and the Graph Attention Network (GAT) [93].

Representing a system of entities as a graph is a powerful way to include complex and heterogeneous prior knowledge into a form amenable to machine learning. Common GNN architectures, however, make nuanced and often unspoken assumptions about the behavior of their input graphs, which are not valid in some learning tasks. We summarize the common GNN assumptions as follows:

- **Edge Uniformity**: Almost all GNNs use a permutation-invariant aggregation scheme to pass information between nodes (e.g., max, mean, or sum functions), which allows for GNN convolutions to be applied to all nodes irregardless of number of edges; however, these methods implicitly assume that any two edges (with the same features if applicable) are equivalent. One notable exception to this assumption is the Graph Attention Network (GAT), which uses edge-specific attention to weight edges during message aggregation [93].
- **Homophily**: Like nodes are more likely to be connected [60]. Graphs in which unlike nodes are often connected are considered *heterophilous*.
- **Locality and Relational Equivalence**: The properties of a node are influenced by their K-th local neighborhood [45], where K is the number of layers in the GNN. A consequence of locality is *relational equivalence*, which posits that any two nodes with the same K-th order relations will have equal GNN representations.

These GNN assumptions are valid in many domains, however, some of these assumptions conflict with the perturbation biology premise that we have described above. The molecular relationships described in the scientific literature are often unannotated and highly contextual in behavior, and therefore the resulting biological graphs are likely to represent relationships that have markedly different and unknown behaviors (e.g., stimulatory vs. inhibitory edges). Given this, it is likely that GNNs that make strong assumptions of *edge uniformity* and *relational equivalence* will not be able to learn distinct edge behaviors due to the dearth of molecular interaction annotations. That said, continued scientific discovery and more detailed categorization of molecular relationships may overcome this limitation in the future and enable the construction of biological graphs suitable for GNNs. Alternatively, some GNN architectures with a weaker assumption of *edge uniformity* and *relational equivalence*, such as graph attention networks [93], may be more appropriate for cellular signaling tasks. The development of novel GNN mechanisms, such as joint learning of edge-specific features during training, may also overcome these limitations.

Recent work has shown that traditional GNN architectures perform poorly on heterophilous graphs [60]. Given our premise of cellular signaling and, assuming that we are using gene expression as the target variable, then biological graphs for this learning task are likely to be largely heterophilous. For example, many molecular interactions directly impact the gene expression (transcription factor regulation, ubiquitation/degredation, miRNA regulation, etc.) while others will indirectly influence the expression (phosphorylation, protein complexes, etc.) via latent (i.e., unmeasured) cell signaling. Regions of the biological network describing protein-protein signaling cascades are likely to be largely heterophilous (signaling cascades), while local regions of gene regulation will be highly homophilous (e.g., transcription factors, GRNs). Given these conditions, it is likely that GNN architectures that make strong assumptions of homophily are likely to perform poorly on cell signaling graphs. While there has been work to develop suitable GNNs for heterophilous graphs [106, 103, 107], it remains an open challenge.

The depth of the GNN, or the number of layers in the network can also prevent challenges in this domain. A GNN operates by subsequent layers of *message passing* between nodes. As a result, the information from any given node can only propagate within the K-th neighborhood defined by the number of layers (K) in the network^4^. For example, a one-layer GNN can only pass information to its immediate neighbors. Deep GNNs (many layers) tend to generate node representations that are very similar to each other, a phenomenon called *oversmoothing*, which can significantly degrade performance [14]. Although many methods have been proposed to address oversmoothing [14, 104], there is no one-size-fits-all approach and remains a limitation in many prediction tasks. In the context of cellular signaling, biological pathways often involve deep signaling cascades and complex gene regulatory networks, and this suggests that deep networks are essential to accurately model the process, and GNNs may therefore suffer from oversmoothing.

## 3 Related Work

### Structural equation models (SEM)

A class of methods that focus on modeling processes with known or postulated dependencies between variables or latent variables. These methods are commonly used in the social sciences to evaluate the appropriateness of causual structure between variables [11]. Many SEMs assume simple linear relationships between variables and struggle to scale to a large number of variables, and this limits their application to many tasks [22].

### BioChemical Reaction Networks (BRN)

A mathematical representation of biological systems that comprise interactions between biochemical species and molecular entities. BRNs can be used to model the behavior of biological systems at various levels of granularity, and modeling strategies include ordinary differential equations (ODE), Boolean Networks, Markov processes, Kinetic rate laws, and agent-based systems. Many BRN methods focus on modeling detailed kinetic systems, and while some of these methods are capable of scaling to a large number of entities, many of them focus on systems with fewer than a thousand parameters [56]. Some general limitations of mechanistic models, such as ODEs, are the difficulty of fitting parameters, especially as the system size scales.

Modeling **signal transduction** is a key goal of systems biology research, and there has been significant work in this area. An early model, called PARADIGM^5^, used factor graphs to model cell signaling [91]. More recently, Boolean networks have been used to create logic models of cellular signaling [89]. Some pathway knowledge bases have included modeling techniques directly into their platform, such as ReactomeFIViz [13] and the breadth and ubiquity of cell signaling modeling highlights the integral role that it has in biological research.

In the domain of cancer drug response, several **mechanistic models** of cell signaling have been proposed to address the scalability issues of BRNs. In 2018, a mechanistic pan-cancer pathway model was introduced, which could learn the ODE parameters for a system of ∼100 proteins/genes and <10*e*4 reactions in approximately a week of training on a system of 400 CPUs [26]. More recently, a machine learning approach termed *CellBox* was proposed to model perturbation biology using an ODE (∼10,000 interactions/*w*_*ij*_) and solved with gradient descent optimization. These methods address a limitation of many machine learning algorithms that lack the interpretability of predictions. ODE models are both transparent (simple, known mathematical model) and traceable (outcomes can be traced along the inferred biological network) [102], and are likely to be more trustworthy and actionable than traditional machine learning methods.

A foundational method that incorporates prior knowledge into deep learning is the **Visible Neural Network (VNN)**. VNNs use prior knowledge to constrain artificial neuron interactions and encourage the model to mimic the behavior of true biological systems [59]. The VNN did this by enforcing a parent-child relationship of Gene Ontology latent states and mapping genotype behavior onto them to predict cellular phenotypes. This work showed that including prior knowledge constraints could aid in the interpretability and usefulness of a model, while maintaining the predictive performance of traditional deep learning approaches. More recent work developed *DrugCell*, which uses a VNN to predict drug response and synergy [46]. A limitation of current VNN methods is that prior knowledge and interpretation are restricted to the ontology that is used. Specifically, DrugCell does not provide molecular-level prior knowledge and assumes hierarchical prior knowledge (i.e., requires directed acyclic graphs).

Another method that aims to incorporate prior knowledge into deep learning is **physics-informed neural networks (PINN)**. These methods use a unique loss function to enforce a set of prior knowledge constraints encoded as partial differential equations (PDEs) [72]. To our knowledge, PINNs have not been applied to systems biology or drug response modeling and tend to focus on domains where the constraints are well known and can be encoded as a system of PDEs.

**Knowledge distillation** was proposed to distill the knowledge and high performance of a large model, or an ensemble of models, into a smaller or single model, thereby reducing inference overhead [36]. A recent paper used a similar method to distill the knowledge from a slow but highly mechanistic cellular model into neural networks. By doing so, they were able to obtain an accurate mechanistic-trained deep learning model, which was orders of magnitude faster at inference than the alternative mechanistic model [94]. This method may represent an alternative means of training robust mechanistic-like deep learning models while maintaining the fast inference. Importantly, this approach is likely to suffer from issues of interpretability and explanability because even though the deep learning model is trained on a data from a mechanistic model it is still a “black box,” which inhibits clear understanding of the internal prediction logic.

### 3.1 Contributions

This work is inspired by SEMs^6^ ability to inject causal structure into modeling strategies and by methods such as GNNs and VNNs that combine heterogeneous prior knowledge with powerful data-driven deep learning methods. To overcome the limitations that prevent the application of current methods to cellular signaling, we present a method termed *Graph Structured Neural Networks* (GSNN), which enables the inclusion of prior knowledge encoded as directed graphs with allowable cycles (e.g., feed-back loops, auto-regulation, etc.). We also introduce the *GSNNExplainer* method, which can be used to inspect GSNN prediction logic in a biologically relevant form. We show that the GSNN algorithm can be used to effectively predict perturbed expression (LINCS L1000) and cell viability for single (PRISM) and combination (NCI Almanac) drug perturbations. Finally, we demonstrate how these methods can be used to prioritize drugs that induce a disease-specific response and evaluate using FDA drug indications.

## 4 Methods

### 4.1 Problem Description

In this project, we present a learning problem in which we are given a graph, *G*, which has *E* edges and *N* nodes. Nodes can be characterized by their in- and out-degree: *input nodes* have an in-degree of zero and nonzero out-degree, *output nodes* have an out-degree of zero and an in-degree of one, and *function nodes* have a non-zero in- and out-degree. Respectively, edges can be characterized similarly, where *input egdes* are edges from an input node, *function edges* are edges from a function node and *output edges* are edges to an output node and have precedence over function edges. The training data, *𝒟*, has node inputs *x* and outputs *y*. Only the input nodes will have nonzero *x* values, while only the output nodes will have nonzero *y* values. Each observation *i* will have unique values for *x*_*i*_, *y*_*i*_. We propose that the graph learning task is to predict *y*_*i*_ using *x*_*i*_ based on the constraints provided by *𝒢*.

Like GNNs, this problem assumes *locality*, that is, the properties of a node should be influenced by its neighborhood; however, there are several key distinctions from the common GNN inspired tasks, specifically:

- This problem requires a single fixed graph, where different observations will have different input/output node features. This can be viewed as a transductive learning task, and does not need to generalize to unseen nodes or novel graph structures.
- The graph structure does not necessarily connect like nodes, and edges can represent a variety of behaviors (no assumption of *homophily* or *relational equivalence*).
- Relations (edges) will have different behaviors and must be inferred from the data (no assumption of *edge uniformity*).

In this work, we focus on graphs that represent cellular signaling networks; however, this problem description is applicable to model a number of other real-world scenarios, which may benefit from the inclusion of heterogeneous forms of inductive bias. Table 1 describes several alternative domains in which the GSNN method could be applied.

**Table 1:** Alternative domains that the GSNN method could applied to.

| Description | Inputs | Outputs | Inductive Bias |
| --- | --- | --- | --- |
| Fluid flow | Initial flow rate | Flow at a different location or time | Geometric constraints (pipes, landscape) |
| Heat conduction | Initial temp. | Temp. at different location or time | Material geometries (shapes, contacts) |
| Electrical circuits | Initial voltages | Voltage at different point or time | Circuit components and connections |
| Causal modeling | Input variables | Output variables | Latent variable interactions |

### 4.2 Graph Structured Neural Network

We present a deep learning method called Graph Structured Neural Networks (GSNN), suitable for modeling biological signaling networks. The GSNN method is initialized using a structural graph (*𝒢*) that describes the molecular entities (nodes) and the interactions between them (edges); *𝒢* defines a set of constraints in the GSNN algorithm by indicating the allowable interactions and expected latent variables (i.e., molecular entities). *Function nodes* (*f*_*n*_) are parameterized by a neural network, where the number of inputs is equal to the in-degree of node *n* in *𝒢* and the number of outputs of the neural network is equal to the out-degree of node *n* in *𝒢*. The number of hidden channels and network layers^7^ of *f*_*n*_ are user-defined hyperparameters, which allow for variable model capacity. We rationalize that proteins with fewer inputs or outputs can be better described by more simple functions and therefore we provide the option to scale the number of hidden channels in *f*_*n*_ based on the in- or out-degree of node *n* .

GSNN layer updates can be performed by masked linear operations, an example of which is shown in panel A of figure 2. Parameters are not shared between *function nodes*, therefore, each function node will learn distinct relationships between input and output edges. The GSNN method operates by evolving the *edge* latent representation via sequential GSNN layers^8^. Each layer will update the latent edge values so that the output edges of a *function node* are predicted from the input edge values of the previous layer. Iterative edge updates allow the information to propagate through the structural graph a path length of *L*, where *L* is the number of layers in the GSNN. The latent features are representative of *edge* state. Note that this aspect is divergent from GNNs where latent representations typically characterize the state of a *node*. This allows the GSNN method to learn nonlinear multivariate relationships between input edges and output edges. As we noted in the Introduction, there is a temporal aspect to cellular signaling such that many molecular entities will have a latency between input and output signals. As a means of modeling this edge latency, we include residual connections at each consecutive layer, which allows for an “accumulation” of signal and provides a mechanism to learn edge latency. Residual connections have also been shown to mitigate gradient vanishing issues that are common in deep networks [32].

**Figure 1:**
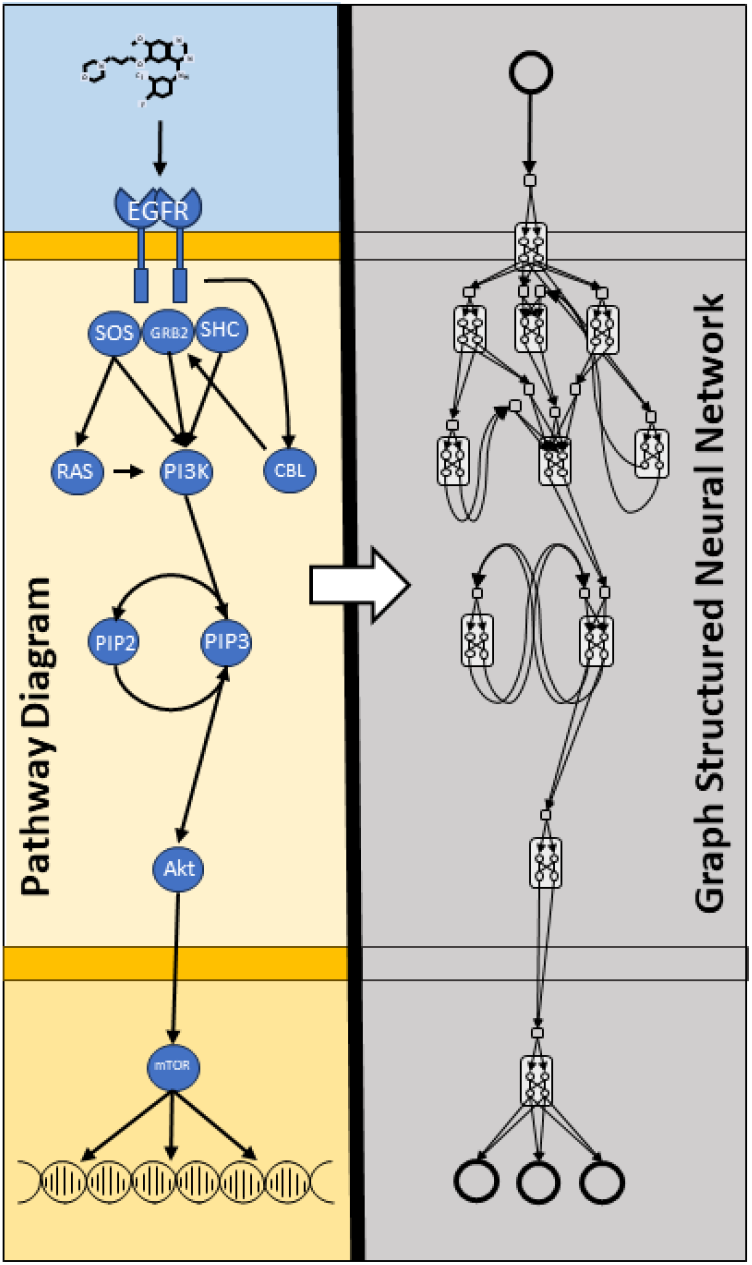
Graph structured neural network (GSNN) summary figure. An example pathway diagram used to describe cellular signaling knowledge (left). A GSNN model built from the pathway diagram. Although not visualized, entity-specific features (e.g., gene mutations, expression, or other ‘omics) can be included as additional input to each entity (right).

**Figure 2:**
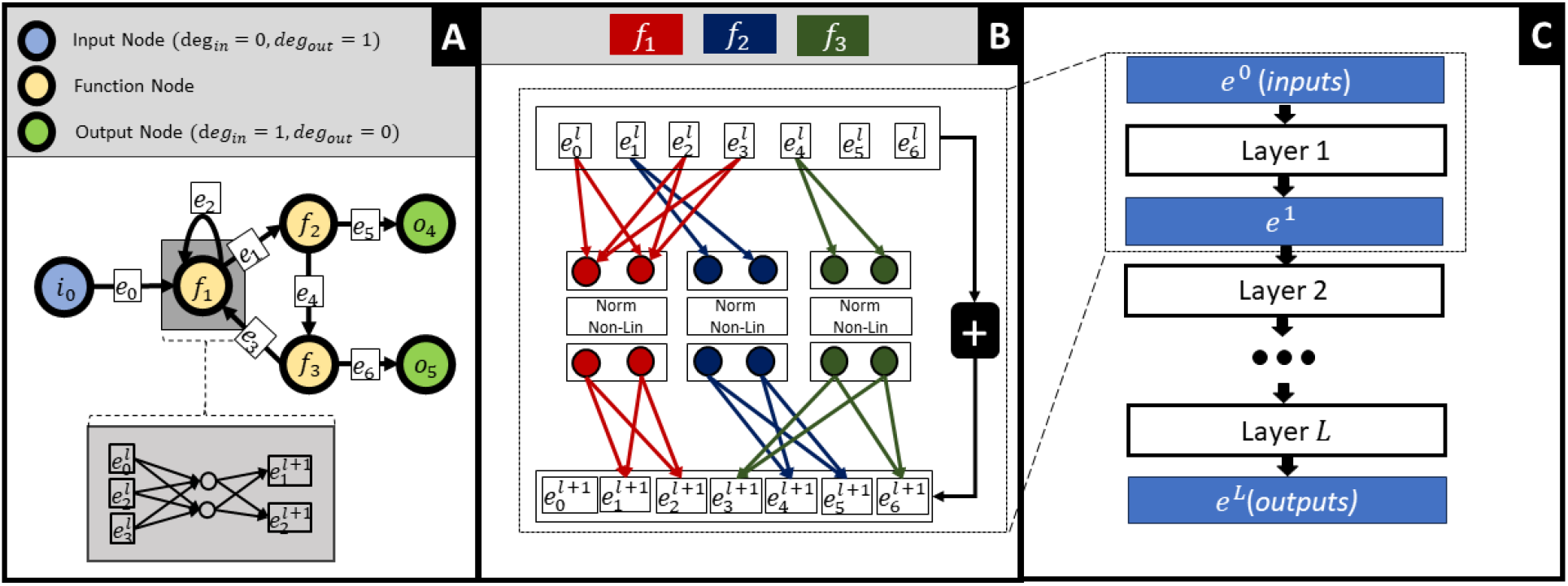
A toy example demonstrating how any given graph structure can be formulated as a feed forward neural network with masked weight matrices. Each yellow node in the left graph represents a fully-connected 1-layer neural network with two hidden channels (Note: function node neural networks can optionally be multi-layer). Panel A describes the structural graph (*𝒢*) which imposes constraints on the GSNN model. Panel B depicts how the edge latent values (*e*_*i*_) can be updated in a single forward pass. Note that panel B shows sparse weight matrices, where the missing edge connections are equal to zero. The plus sign in panel B indicates a skip connection from the previous layer.

**Figure 3:**
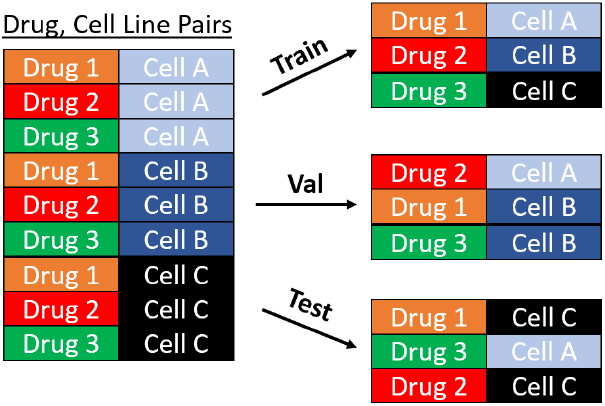
The data partitioning scheme used to train and evaluate the GSNN algorithm.

**Figure 4:**
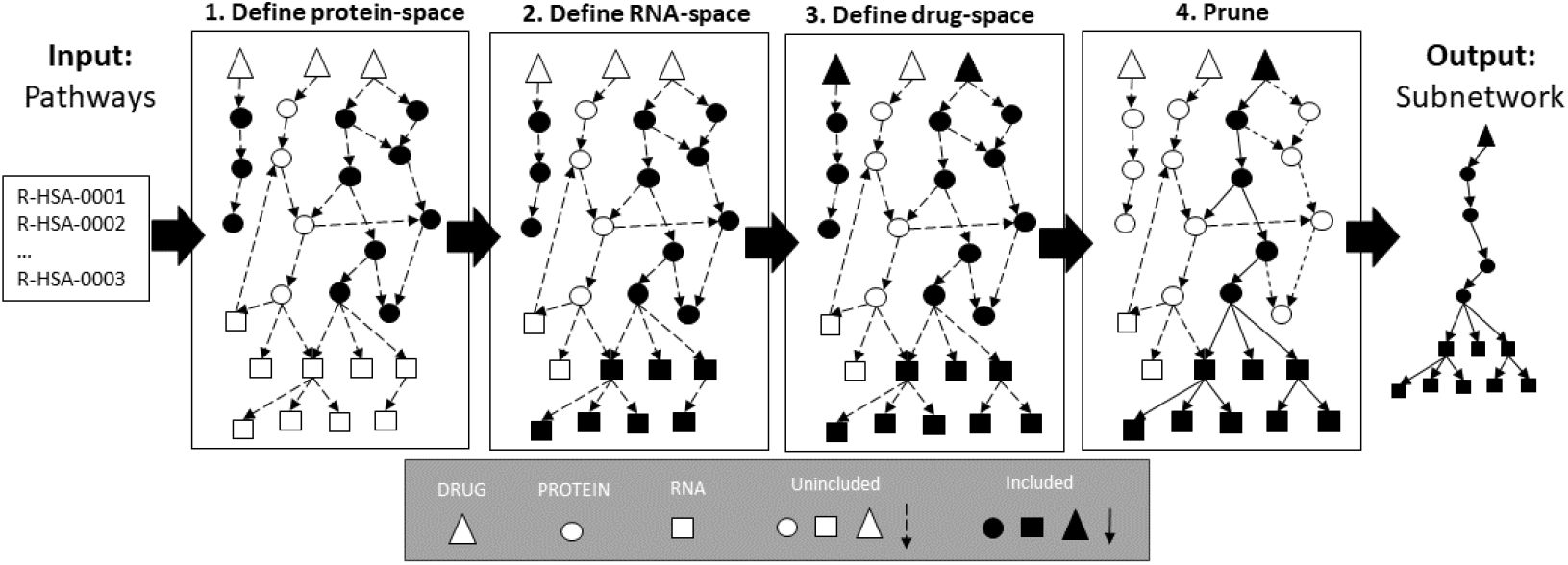
Network construction diagram. Given a full biological network relating drugs, proteins and RNA (mRNA + miRNA) we construct a subnetwork focused on user-provided pathways. Each panel describes a sequential step in the construction. First, we use a user-provided set of pathways to define a *protein-space* that includes all proteins from the full network that have membership in at least one input pathway. The second step defines the *RNA-space* as all RNAs that are regulated by at least one protein in the protein-space; entities from multi-step regulation (e.g., TF->miRNA->mRNA) are also included. The third step is to define the *drug-space* as all drugs that target at least one included protein. Finally, there is a pruning step that removes any nodes that do not regulate a given proportion of downstream RNA outputs (user-defined; typically ∼ 25%). The output of this process is a biological subnetwork that comprise the molecular entities and relationships that describe the input pathways.

**Figure 5:**
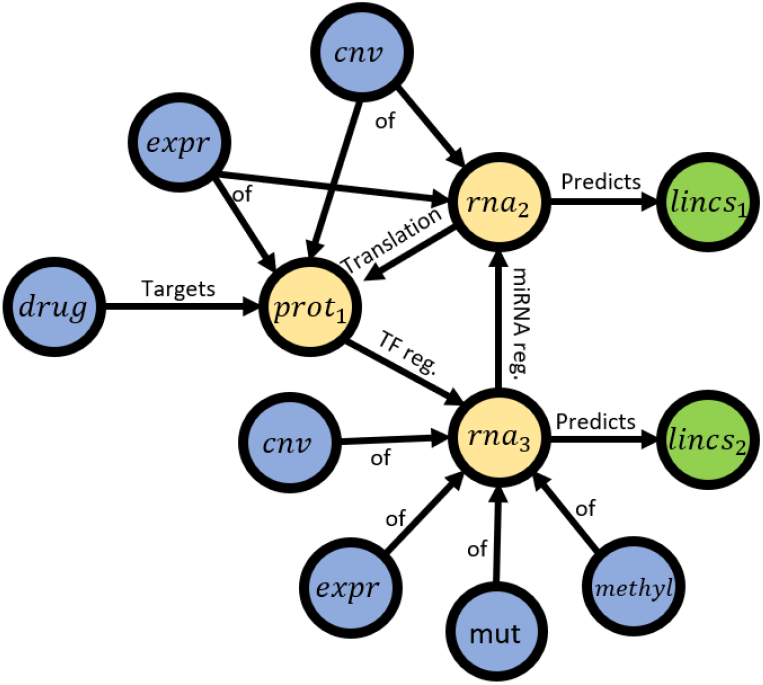
A toy example demonstrating how a biological network for perturbation modeling can be created. Input nodes include both drug and ‘omics features (blue), output nodes represent LINCS measurements of a respective RNA molecule (green), and proteins and RNA molecules are function nodes (yellow). Note that while we have added edge labels in order to demonstrate what a biological network can represent, we do not include edge-specific features in the GSNN modeling.

To efficiently implement the GSNN method, we conceptualize the edge-updates as a series of masked linear layers. The weight matrices have dimensions (*E, N* ∗ *C*), where *E* is the number of edges in *𝒢, N* is the number of function nodes in *𝒢*, and *C* is the number of hidden channels in each *function node*. Implementing dense matrix multiplications of these linear layers would require undesirable memory and compute resources, making this method applicable only to relatively small graphs. Fortunately, we can use sparse matrices, which massively reduces the required memory.

The GSNN method can be considered a residual network [32], such that *x*_*l*+1_ = *F* (*x*_*l*_) + *x*_*l*_ with the added constraints imposed by the structural graph *G*. Function node parameters may be optionally shared across layers, which prevents the model parameters from scaling with depth; however, in practice we find that the GSNN model performs better when parameters are not shared across layers. We also optionally add self-edges to all function nodes to allow them to incorporate self-information from the previous layer. Taking lessons learned from traditional residual networks, we also include normalization layers. The ResNet model uses batch normalization [32], however, due to the memory requirements of the GSNN method, we are required to use small batch sizes for training, and therefore batch normalization is unlikely to perform well. Instead, we use layer normalization [5] *within* each function node to prevent data leak between function-nodes.

We include options to use Kaiming/He or Xavier/Glorot weight initialization [33, 49]; however, since the function nodes can only access a subset of inputs and outputs, we use the in-degree 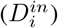 of node *i* in the input graph *𝒢* as the “fan in” value and out-degree 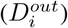 as the “fan out” value. With this modification, our weight initialization methods are described by:

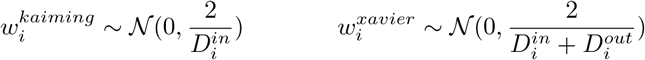

We implement the GSNN model in Pytorch [68] and, at the time of writing this, Pytorch’s native sparse matrix multiplication is not well optimized for batched operations. To improve on this, we use the Pytorch Geometric package [24] to perform mini-batching and formulate our sparse matrix multiplication as a Pytorch Geometric graph convolution. This approach is markedly faster, particularly when operating on a GPU, than using Pytorch’s native sparse matrix multiplication.

### 4.3 Model Evaluation

To evaluate performance, we use Monte Carlo Cross-Validation (MCCV) [100] to randomly subsample train (60%), validation (20%) and test (20%) data subsets. We run multiple folds (*n* ≥ 3) and choose the best performing model within each fold using the validation set. We report performance as the average Pearson correlation 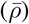 evaluated in the test set across all folds. Each data partition is a disjoint set of *(drug, cell-line)* pairs (e.g., Imatinib, A549). This evaluation approach allows the model to train on all drugs and cell lines while evaluating on unseen drugs and cell line pairs. Note that this partitioning scheme evaluates the performance of the model within the measured drugs and cell lines, but generalization to unseen cell lines is not evaluated. A model evaluated with this approach can be effectively used to impute missing drug and cell line combinations.

To measure the performance of the model, we use the mean Pearson correlation 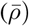 [25] across all predicted LINCS gene outputs (*y*_*i*_), where *N* is the number of LINCS gene outputs.

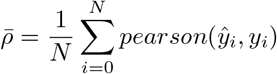

To determine significant performance differences between models, we perform two-sided paired t-test, adjusted for the number of tests (n=3) with the Bonferroni correction [31].

### 4.4 GSNN performance on random networks

As we have discussed, the *no free lunch theorem* suggests that including prior knowledge in deep learning algorithms has the potential to improve prediction performance compared to models without prior knowledge. We have developed the GSNN method to do precisely this, and expect that if the GSNN algorithm outperforms the baseline, then the prediction advantage is due to inclusion of prior knowledge; however, it is possible (albeit unlikely) that the GSNN performance could be independent of the prior knowledge (i.e., due to a different aspect of the GSNN algorithm). To test this assumption, we compare the performance of matched hyperparameter GSNN models that are initialized using either 1) a randomized biological network or 2) the true biological network. If the GSNN performance is due to the inclusion of accurate prior knowledge, then we should see significant performance differences between models trained on true and random prior knowledge.

*Randomization* of the biological network is performed by sampling new edges of the input graph (*𝒢*). This approach maintains the total number of edges, but not the in- or out-degree of function nodes. *Input edges, function edges* and *output edges* are randomized independently to maintain the same number of input and output edges. For all *input edges*, only the destination of each edge is randomized, and similarly for *output edges* only the source of each edge is randomized.

### 4.5 Biological Graph Construction

To create a biological network suitable for modeling cellular signal transduction, we used the resources listed in Table 2. In our biological network, allowable input nodes include: DRUG, EXPR (rna expression), CNV (copy number variation), MUT (mutation) and METHYL (methylation input). We may refer to EXPR, CNV, MUT or METHYL as OMIC nodes and these input represent cellular context (e.g., cell line encoding). All the PROTEIN and RNA nodes will be *function* nodes. In this project, all output nodes refer to a LINCS L1000 gene and will have a single edge connection from the respective RNA node (e.g., RNA P53 → LINCS P53). Note that we do not include DNA molecules as separate entities; rather, we collapse this representation within the RNA nodes. For example, a transcription factor that targets a DNA gene will be encoded as targeting the respective RNA gene. In the current formulation, including DNA nodes offers little advantage and would increase the computational complexity of the model during training and inference.

**Table 2:** Literature curated resources documenting molecular interactions.

| Resource Type | Resource | Expert reviewed | Source Node | Target Node | Refs |
| --- | --- | --- | --- | --- | --- |
| Drug-Target | The Cancer Targetome | Some | Drug | Protein | [12] |
| Drug-Target | CLUE Repurposing | No | Drug | Protein | [20] |
| Drug-Target | STITCH | No | Drug | Protein | [47] |
| Protein-protein interaction | Omnipath | Some | Protein | Protein | [90] |
| Transcription Factor Gene Regulation | Omnipath (Dorothea) | Some | Protein | RNA | [90, 27] |
| miRNA Gene Regulation | Omnipath (MiRNA) | Some | RNA | RNA | [90] |
| Translation | NA | NA | RNA | Protein | NA |

The resources listed in Table 2 span many pathways and processes, of which many are likely to be irrelevant to gene regulation or common chemical perturbation pathways. Furthermore, the number of trainable parameters in the GSNN model scales with the input graph size. For these two reasons, we developed a process to select a subset of proteins that are likely to be predictive of chemical perturbation signaling processes. This graph construction process relies on a set of user-defined Reactome pathways. We then define the intersection of pathway proteins as the *protein-space* of *𝒢*. Next, we define the *rna-space* as all RNAs that are targeted by a member of the *protein-space*; Optionally, the user can provide a depth to which RNA descendents will be included, which allows for inclusion of more complex gene regulation networks (e.g., protein->miRNA->mRNA). Any drugs that have a reported protein target in the *protein-space* are then added to the *drug-space*. The *LINCS-space* is the intersection of available LINCS genes and *rna-space*. We then include all edges from the resources listed in 2 if both source and destination are in one of the node-spaces (the user can optionally choose a subset of resources to include). Next, there is a drug pruning step based on how many LINCS nodes are descendants (the default is 25% of the number of LINCS nodes). This pruning step is used to remove drugs that are overconstrained by the available prior knowledge and therefore cannot impact the prediction of most outputs. PROTEIN and RNA nodes are also pruned if they do not have downstream LINCS genes. OMICS nodes are added based on the intersection of available ‘omic features’ and protein-space or rna-space. In this construction process, an OMIC node (e.g., EXPR P53) can have edges to a PROTEIN, RNA, or both (example: see *prot*_1_ and *rna*_2_ in Figure 2). For instance, we rationalize that RNA expression is relevant to both RNA and protein behavior, and therefore we make the respective OMICS easily accessible to both.

Protein inclusion based on pathways makes the assumption that all output nodes (LINCS expression) in the graph can be well predicted via the entities we include; however, this assumption is likely violated in many cases, especially for small pathways. Including larger or multiple pathways could mitigate this issue, allowing for cross-talk between pathways. The exclusion of a drug’s targets is likely to prevent the GSNN method from effectively predicting the response to that drug. Similarly, exclusion of critical transcriptional regulators of an RNA molecule may lead to worse performance on the respective LINCS output. Therefore, we manually select pathways to balance the graph size (so that the GSNN method is memory- and compute-efficient), while including molecular entities relevant to drug-response.

### 4.6 Baseline Models

We compare the performance of the GSNN algorithm against several baseline algorithms: Artificial neural network (NN) [62], Graph Convolutional Network (GCN) [45], Graph Attention Network (GAT) [93] and Graph Isomorphism Network (GIN) [98]. We also compare performance against a “cell-agnostic” Neural Network, in which all cell-context specific features are removed (e.g., expression, mutation, methylation, and copy-number variation features are removed from the input). This provides a baseline that characterizes the prediction of the average response of each drug, across all cell lines.

We implement the NN baseline using two layers with the Exponential Linear Unit (ELU) activation function [18] and batch normalization [41]. All Graph Neural Networks (GCN, GAT, GIN) are implemented with batch normalization layers, jumping knowledge [99], and use the ELU activation function.

### 4.7 Hyper-parameter Tuning

For all three models, we perform a limited hyperparameter tuning within each MCCV fold, and the tested parameters are reported in Table 3.

**Table 3:** Hyper-parameter grid search performed in each experiment for the five tested algorithms. The “Other” line indicates additional Boolean flags that describe specific algorithm behavior.

| Hyper-Param. | GSNN | NN | GIN | GAT | GCN |
| --- | --- | --- | --- | --- | --- |
| Channels | 10, 20 | 100, 500, 1000 | 64, 128 | 64, 128 | 64,128 |
| Layers | 10, 20 | 2 | 5, 10 | 5, 10 | 5,10 |
| Dropout | 0, 0.25 | 0, 0.25, 0.5 | 0 | 0 | 0 |
| Learning Rate | 1e-2, 1e-3 | 1e-2, 1e-3 | 1e-2, 1e-3 | 1e-2, 1e-3 | 1e-2, 1e-3 |
| Other | add_self_edges, scale_channels_by_degree |  |  |  |  |

### 4.8 Drug Dose Transformation

To encode the presence of a drug, each drug node is assigned a scalar value that represents the concentration of that drug in an observation. We transform the drug concentration (*μM*) using the following function:

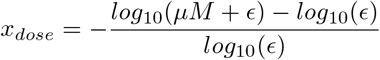

This transformation ensures that the drug effect is log-linear in the relevant therapeutic concentration range, and which can be shifted by choice of *ϵ*. It also maintains that a concentration of 0*μM* (i.e., no drug) will still be equal to zero after transformation and a concentration of 1*μ*M (a typically large dose) is equal to one after transformation. Most of the LINCS L1000 observations were measured with concentrations between -2.5 and 2.5 (*log*_10_(*μM*)), and choosing an epsilon of 1*e* − 6 ensures a logarithmic linear relationship in the concentration range relevant to the LINCS L1000 dataset.

### 4.9 GSNN Explanation: Edge Importance Scores

An advantage of the GSNN algorithm is that we can use the network structure to explain predictions in a form amenable to biological interpretation. Previous work has presented a method for explaining GNN predictions, termed *GNNExplainer*, which identifies a subset of nodes and features that are most involved in the prediction of a given observation [101]. We implement a similar method to explain a given observation predicted by the GSNN network; however, rather than identifying a subset of nodes or features, we identify a subset of *edges*. Given an observation (*x*), baseline observation (*x*_*b*_), and a trained model (*f*_*GSNN*_), we initialize an edge mask (*M*) and use gradient descent to identify a subset of edges that result in comparable model predictions. Further details are available in Algorithm 1. The output of the *GSNNExplainer* method is a subgraph, which includes the most critical or involved edges to predict a given observation compared to a baseline prediction. A key distinction between the *GSNNExplainer* and the *GNNExplainer* is our use of a baseline prediction. GNNs traditionally use a permutation invariant aggregation scheme such as *mean, max* or *sum*, and as such, removing edges or nodes in the GNN setting is analogous to setting the respective latent state to zero; however, in the GSNN algorithm, edges may have some level of endogenous latent activations (perhaps related to ‘omic inputs), even in the absence of drug inputs, and this means that setting an edge latent state to zero is not necessarily representative of “removing” an edge. In the GSNN algorithm, setting an edge latent value to zero is likely to be more analogous to setting the edge to the “average” signaling behavior of the training dataset, which is unlikely to represent any specific observation. To mitigate this challenge, we use a *baseline* observation to characterize the baseline latent edge values. For example, to explain the prediction of a given drug *Drug* = *A*, concentration *Conc* = *c* in a cell line *Line* = *l*, we can use a baseline observation without any drug (*Drug* = *A, Conc* = 0, *Line* = *l*). Using such a baseline allows the *GSNNExplainer* to ignore any endogenous latent activation that may be due to cell type and instead focuses on the edges involved in drug response prediction. Alternatively, one could ask the question *What are the key edges involved in the prediction differences between two different cell lines?* To do this, we can use one cell line as the baseline observation and the other as the input observation, while keeping the drug concentrations the same for both cell lines.

#### Algorithm 1

GSNNExplainer

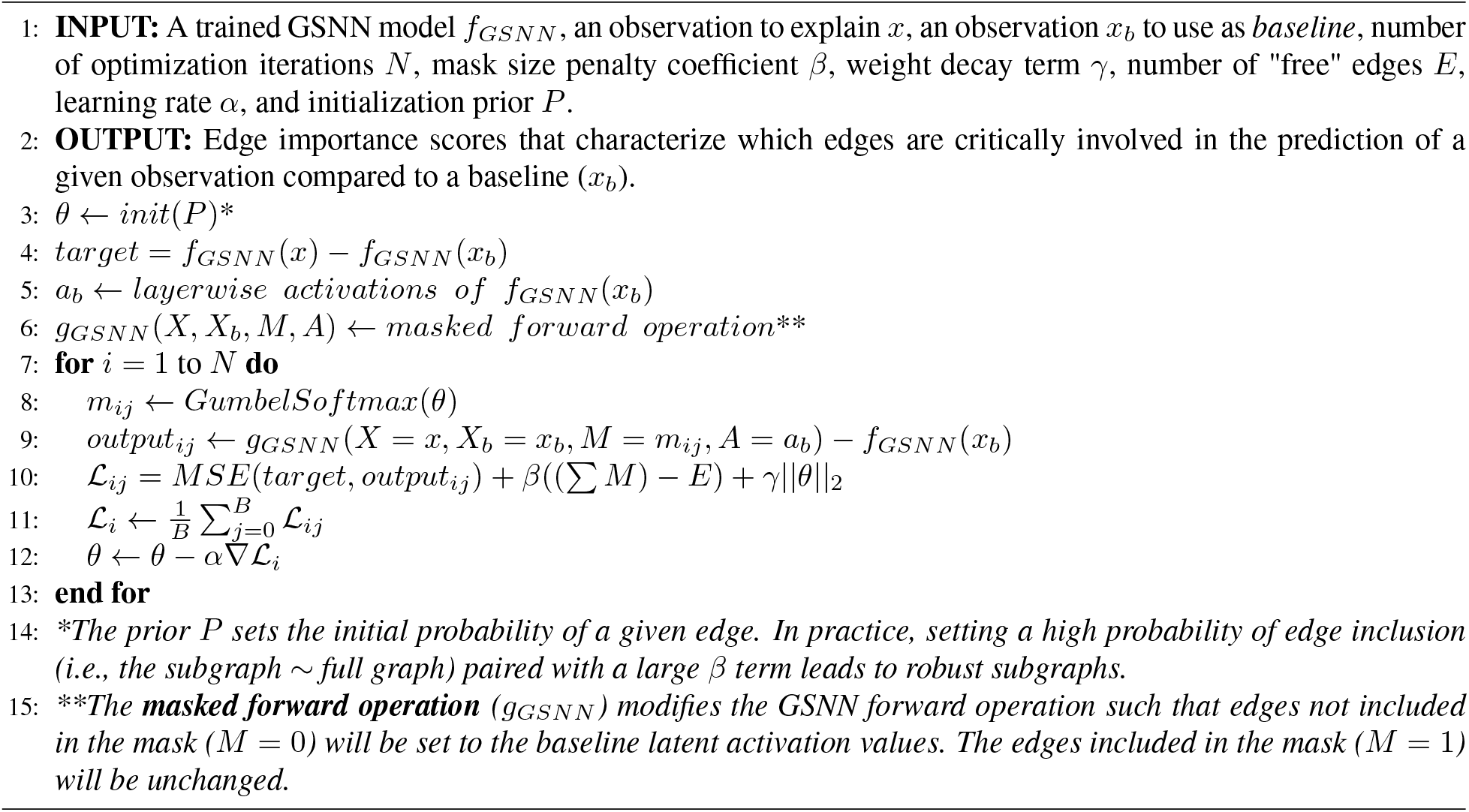

There are a few additional divergences from the methods presented in *GNNExplainer*: First, to induce discrete sampling of edges, we use the softmax-gumbel distribution and anneal the temperature parameter during optimization [61, 42]. Second, we use the mean-squared error (MSE) loss function, which is convenient for multi-output regression. In practice, we initialize the weight mask parameters such that each edge is likely to be selected, thus the optimization starts with almost all edges included in the mask. A strong mask penalty term (*β*) encourages the removal of uninvolved edges during optimization. We also include an optional weight decay coefficient (*γ*), intended to encourage exploration by preventing large (confident) edge parameters. In summary, *GSNNExplainer* produces a biologically relevant explanation of an observation. Using this method, we can interrogate which molecular entities and entity interactions are important for the prediction of an observation.

### 4.10 Drug Prioritization

One of our primary research goals is to use the GSNN method for effective and interpretable prioritization of drugs for nuanced research goals. We base our prioritization procedure on the premise that drugs which induce the same response in all cell lines are unlikely to be good therapeutic candidates due to the detrimental effects on normal cell types. Rather, we seek to identify drugs that induce a selective response in a subset of user-designated cell lines representative of their research goals. For example, researchers may seek a selective response in cell lines derived from a certain primary disease (e.g., breast cancer), or cell lines with shared mutations (e.g., TP53) or an expression pattern (e.g., HER2+).

First, a user must define the contextual response of an ideal drug candidate. This can be done by designating in which cell lines a candidate drug should cause a *desirable* response, termed the “target” lines (*T*), and the cell lines that should have a less *desirable* response, termed the “background” lines (*B*). For example, to prioritize drugs that have a selectively desirable response in breast cancer cell lines, we can assign all breast cancer derived cell lines to the “target” set and all other cell lines to the “background” set:

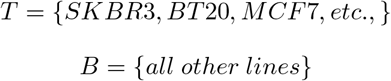

Next, we must choose a metric to quantify the *desirable* response. A desirable response is often measured in terms of cell viability^9^(the proportion of cells alive after the treatment of a drug), where a low cell viability indicates a desirable response. Although cell viability is a convenient measurement for cancer drug response, there are alternative metrics that can be tailored to more specific research questions. For instance, metrics derived from gene expression signatures or single gene expression values may be more appropriate for certain research goals, such as quantifying DNA damage or the activation of apoptotic pathways. For simplicity, however, we chose to quantify the “sensitivity” of a response using cell viability such that low cell viability indicates sensitivity to a drug. We train a probabilistic cell viability predictor network (*f*_*viab*_) using the output of a trained GSNN, such that:

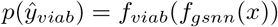

The GSNN parameters are frozen so that they do not change during the optimization of *f*_*viab*_. We use deep ensembles as described by Blundell et al. [50] to quantify the uncertainty in the outcome variable and assume that cell viability is generated from a Beta distribution. Cell viability values are clipped between 0 and 1. Individual viability networks predict two concentration parameters (*a, b*), which characterize the predicted cell viability probability distribution: *P* (*ŷ*) = *Beta*(*a, b*). We parameterize *f*_*viab*_ by a 1-layer neural network and optimize the parameters using the negative log-likelihood and the PRISM dataset [21], which characterizes cell viability after drug treatment in a range of doses. We include all (drug, cell) PRISM observations for which the drug is included in the GSNN model and the cell is included in our GSNN training data. We maintain the same validation and test partitions as used for GSNN training.

To prioritize drugs that create a selective cytotoxic response, we compute the probability that sensitive lines have lower cell viability than resistant lines. To do this, we define the “target” cell viability probability (*P*_*T*_ (*ŷ*_*viab*_)) as a mixture over the “target” lines. Respectively, the cell viability probability of the “background” cell lines is defined as a mixture over all members within the background set (*P*_*B*_(*ŷ*_*viab*_)). We then compute the difference in cell viability between the “target” and “background” lines. Finally, we compute the probability that the target cell lines have an average cell viability lower than the average cell viability of the resistant lines (*p*_*sens*_).

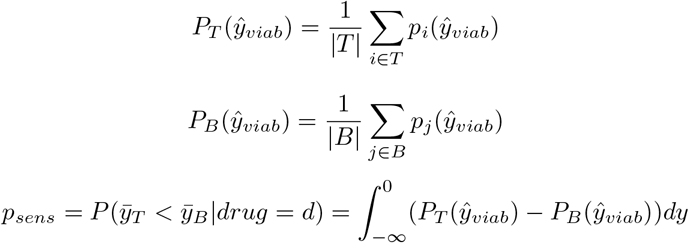

The drugs are then ranked by *p*_*sens*_ to produce a prioritized list such that the top candidates are the most likely to be selectively responsive as defined by the “target” and “background” cell lines. Recognizably, an analytical solution for a mixture of Beta distributions is not convenient, so instead we use Monte Carlo simulations [74] to approximate *p*_*sens*_.

To evaluate the effectiveness of our drug prioritization method, we use disease indications provided by the Drug Repurposing Hub dataset [20]. This dataset provides limited drug annotations that specify the disease(s) for which a drug has FDA approval. To evaluate the rationality of our prioritization algorithm, we generate drug rankings (as described above) where the “target” set includes all cell lines derived from a certain primary disease, termed *target disease*, and the “background” is a set of all cell lines derived from *background disease*. Our expectation is that drugs with FDA indications for the *target disease* will be prioritized over drugs with indication *background disease*. Drugs with indication for multiple disease are not considered. We then quantify the prioritization results using the area under the receiver operator curve (AUROC) such that drugs with an indication for *target disease* are assigned a label of 1 and drugs with an indication for *background disease* are assigned a label of 0. A perfect ranking (AUROC = 1) would be achieved if all drugs with indication for *target disease* are ranked higher than drugs with indication for *background disease*.

## 5 Results

We apply our method to model cellular signal transduction using literature-curated prior knowledge to create a biological network and optimize model parameters using the LINCS L1000 dataset. The LINCS dataset characterizes cell line RNA expression changes in response to chemical perturbations. The L1000 assay measures 978 genes directly and then uses these *landmark* genes to infer the RNA expression of approximately 12 thousand more genes (the “best inferred” and “inferred” features spaces) [85]. We construct a biological network relating proteins, drugs and RNA using the Omnipath resource [90], CLUE compound information [20], the Cancer Targetome [12] and the STITCH database [47].

To limit memory requirements and focus on relevant signaling entities, we generate biological subgraphs that pertain to specific biological pathways. These subgraphs allow us to evaluate the performance of our method on several distinct pathways and data subsets. We define an “experiment” as a choice of biological pathway and network construction hyperparameters. The experiment parameter choices will define which biological entities are included in the biological network, as well as the cell lines, drugs, and observations that are applicable to the respective experiment. In addition, each experiment will randomly assign train, test, and validation partitions. To mitigate the variance in performance due to partition sampling, we run each experiment in replicate (*n* = 3), such that each replicate will share all hyperparameters, but will have unique data splits and weight initialization. Within each replicated experiment, the validation set is used to select the best performing model from a hyperparameter grid search, and we report the average test performance in Table 4b. Table 4a describes the experiment hyperparameters and lists the “primary pathways” that were used in network construction; for further details regarding the experiment construction parameters, see Supplemental section 9.1. Table 4c reports the results of a paired t-test comparing the performance of GSNN and NN (the two algorithms with the highest performance). The GSNN algorithm obtains the highest mean Pearson correlation in all three experiments and significantly outperforms (Family-wise error rate (FWER) < 0.05) the NN in experiment 1. Of the GNN algorithms, the GIN obtains the highest mean Pearson correlation but still underperforms compared to the NN and GSNN algorithms.

**Table 4:**
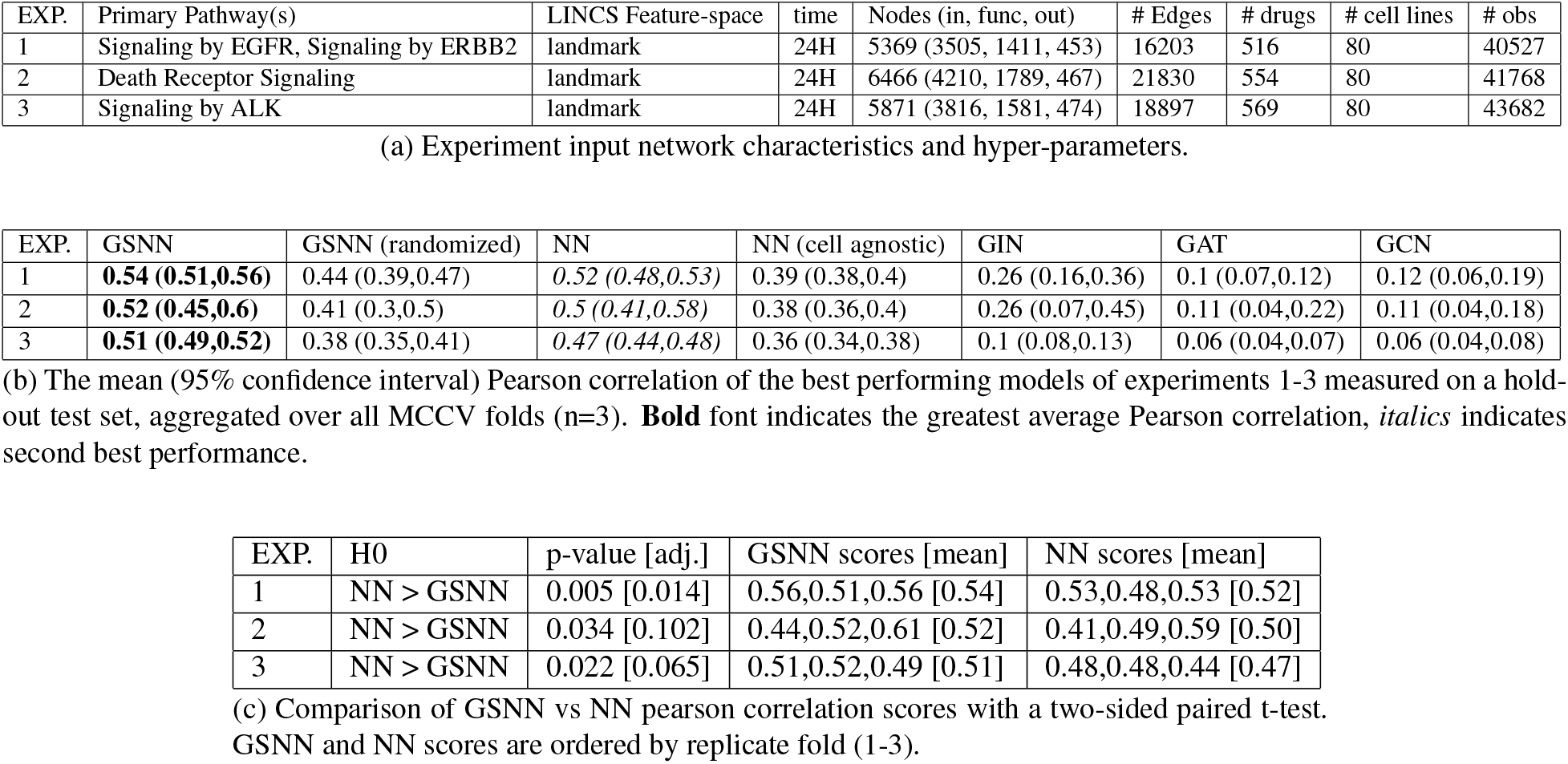
Algorithm performance on LINCS L1000 perturbation data.

### 5.1 Local Performance Advantages

Perturbation biology requires the prediction of many outputs based on numerous inputs. Primary sources of variance in response to a perturbation include drug, concentration, and cellular context. In such a complex system, it can be useful to inspect the *local* performance of a predictive model, which we define as the performance grouped by attribute or performance within a subset of outputs. *Local performance* can help investigate questions such as: *Which drug(s) perform best? Which genes (outputs) are well predicted?* Investigating such model behavior can help highlight where the GSNN method works well and where it falls short.

The GSNN model is constrained by the biological network (*𝒢*) constructed from literature-curated datasets. It is therefore likely that the biological network will have quality biases toward well-studied pathways or relevance to certain cellular contexts (i.e., the contexts that are most commonly studied). Additionally, the choice of network construction hyperparameters (particularly the choice of pathways) may benefit the prediction of a subset of drugs or genes. A plausible outcome is that such biases will translate to local regions of the biological network that are more useful or accurate in predicting the expression change of particular genes. Alternatively, certain drugs, cell lines, or observations may be particularly well predicted because of the GSNN inductive bias. To investigate this, we examine the *local* performance advantages of the GSNN model compared to the NN (top row of Figure 6) and to the GSNN model initialized with random prior knowledge (“GSNN-rand”; bottom row of Figure 6). We use the best model from each fold (N=3) to evaluate performance grouped by various attributes. For example, 6a shows the results of the observations grouped by drug; all observations with a given non-zero drug concentration are grouped and the performance is computed as an average across all outputs and experiment replicates. We evaluate group-performance in the test set and test the statistical significance of each group comparison using a paired two-sided t-test and adjusted for multiple tests using the Benjamini-Yekutieli or Benjamini-Hochberg^10^ false discovery rate (FDR) method [10, 9] and an FDR threshold of 0.1.

**Figure 6:**
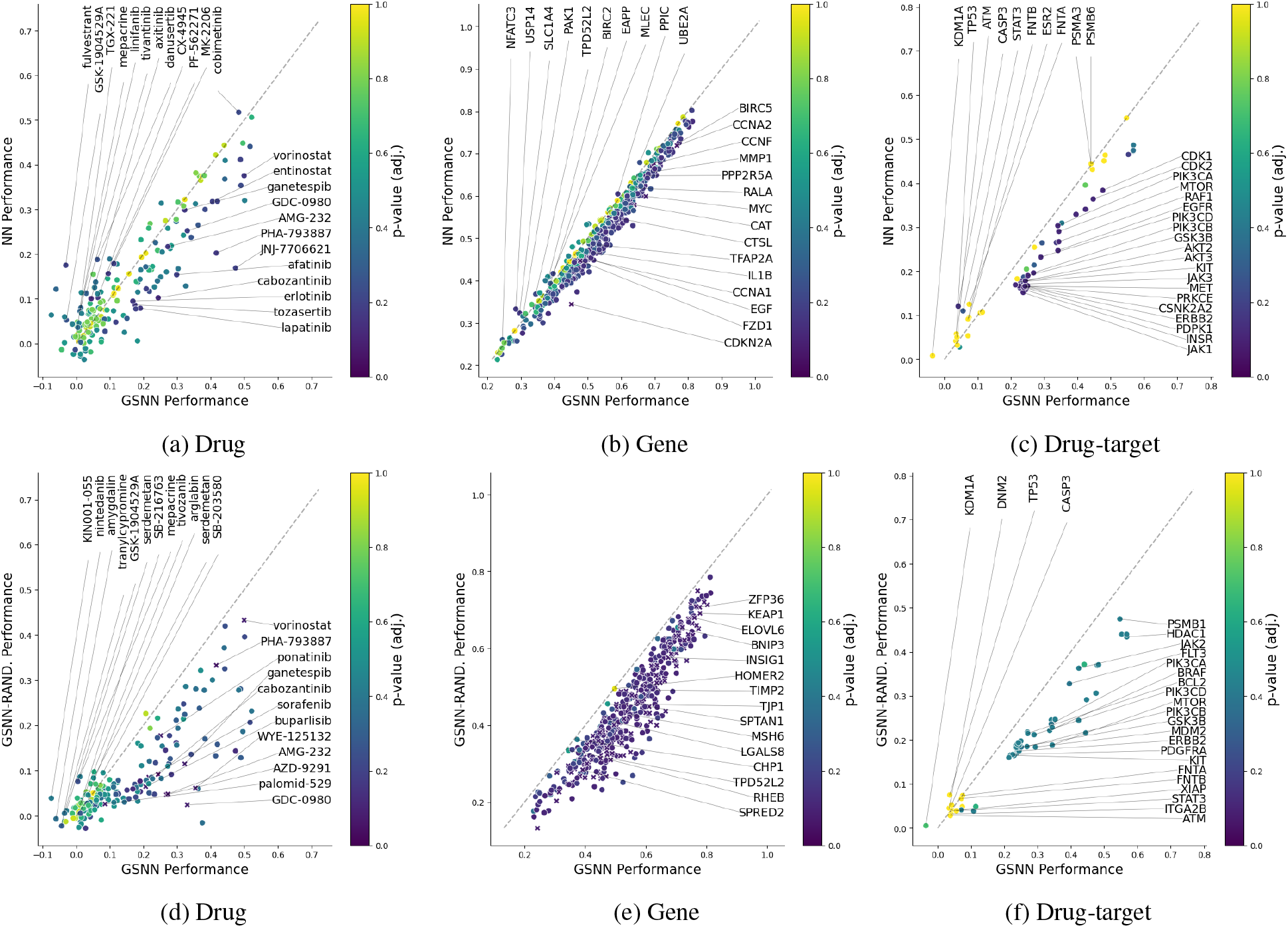
Test performance comparison between the GSNN model (x-axis) and either the NN (top row; y-axis) or Randomized-GSNN (bottom row; y-axis) when grouped by several attributes (Drug (left), gene/output (middle) or drug-target (right)). Performance is reported as the Pearson correlation averaged over predictions from the best model from each MCCV fold of Exp-1 (EGFR + ERBB2 signaling). The gray dashed line on all plots represents equivalent average performance across folds; any groups lying under the dashed line are better predicted by the GSNN algorithm (i.e., “GSNN performance advantage”). Significance was determined using a paired t-test (n=3) and groups with a p-value less than 0.1 are denoted with an ‘x’. Groups with fewer than five observations in each fold (n=3) are not included. Note that we have limited power to detect significant performance differences because we only ran three replicates.

The drug targets with the most significant prediction advantage are proteins that are integrally involved in the pathways chosen for Exp. 1 including EGFR, ERBB2 and CDKs. The two most significant drug-targets with prediction advantage by the NN are CASP3 and ATM. Furthermore, there are many drugs and drug-targets that are poorly predicted by both the GSNN and NN algorithms (r < 0.2), which may suggest poor data quality (e.g., noisy L1000 measurements) or low data volume. There are also several drugs that are predicted quite well (r>0.5) by both GSNN and NN, and this may suggest that these drugs induce a simple response (i.e., “easy” to predict) or have high data volume (many measurements for the given drug).

### 5.2 GSNN Performance on random networks

The premise of our research assumes that prior knowledge encoded in the biological graph will be useful for the prediction of the endogenous variable. To test this assumption, we compare the performance of the GSNN algorithm when using true or randomized biological graphs. We find that GSNN models that use the true biological graph result in an average performance improvement of 24% (95% CI: 11.4% - 40.1%) compared to hyperparameter-matched GSNN models that use a randomized biological graph. Figure 7 shows the performance advantage when using the true biological network. Randomization of the biological network decreases performance in all three experiments. Interestingly, the randomized GSNN model outperforms the cell-agnostic NN and all three GNN algorithms, suggesting that the GSNN model is capable of accurate predictions despite random inductive bias. Of note, the biological graph prior knowledge that the GSNN utilizes has two forms of inductive bias, 1) the interactions between molecular entities and 2) the molecular entities themselves. Our randomization scheme can only remove inductive bias from molecular interactions and there may still be predictive value in the knowledge of molecular entities themselves.

**Figure 7:**
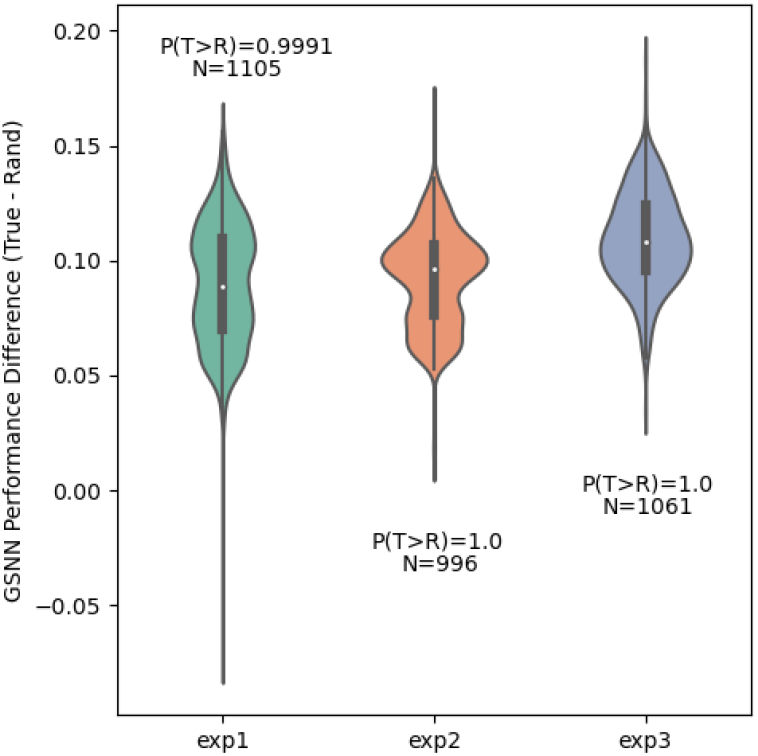
The performance differences between hyper-parameter and replicate matched GSNN-true (true bio. network) and GSNN-rand (random bio. network). Performance evaluated with Pearson correlation and differences computed by Δ*r* = *r*_*GSSN*−*true*_ − *r*_*GSNN*−*rand*_. P(T>R) is calculated as the proportion of positive differences (i.e., proportion of GSNN-true performances that are greater than GSNN-rand performances).

### 5.3 Explaining Predictions using Edge Importance Scores

Significant progress has been made toward the interpretation of traditional neural networks, including attribution methods [58, 86] and black-box explanation methods [73]; however, applying these methods to cellular signaling models does not express model prediction logic in a biologically relevant form. This limitation is exacerbated in traditional neural networks, since the prediction logic is unlikely to accurately capture the sequence of molecular interactions and, therefore, even accurate explanations of traditional neural network logic are not necessarily useful to understand the underlying biology. The network-based architecture of the GSNN presents new approaches to interpret explanations in a way that may be more useful to biologists. To do this, we implement a method termed *GSNNExplainer*, inspired by previous work [101], which explains an observation by identifying a subset of edges that are “important” to the prediction of the model.

Subgraph explanations produced by the *GSNNExplainer* can be conceptualized as a testable hypothesis of the true underlying drug response. For instance, the subgraph shown in Figure 8c highlights the key molecular entities that the Exp. 1 GSNN model uses for the prediction of outcome variables. Subsequent work could validate these explanations by comparison with the literature or by performing wet-lab bench testing to confirm the involvement of molecular entities. For example, Figure 8c clearly shows ESR1 (Estrogen Receptor) has an important role in the prediction of the response to BRD-K50691590 (proteasome inhibitor). To confirm or deny the involvement of this transcription factor, researchers could use experimental assays, such as chromatin immunoprecipitation sequencing (ChIP-seq) [67], to investigate the activity of ESR1 (or other implicated transcription factors) in the response to BRD-K50691590. Representing drug response explanations as a testable hypothesis can be used to build knowledge (e.g., identify important molecular entities) or to encourage user trust in the model (through validation of explanations). Although traditional neural networks can explain predictions using methods such as SHAP [58], explanations can only relate inputs and outputs and lack a representation of intermediate molecular entities and therefore can be challenging to use as a testable hypothesis of the underlying biology.

**Figure 8:**
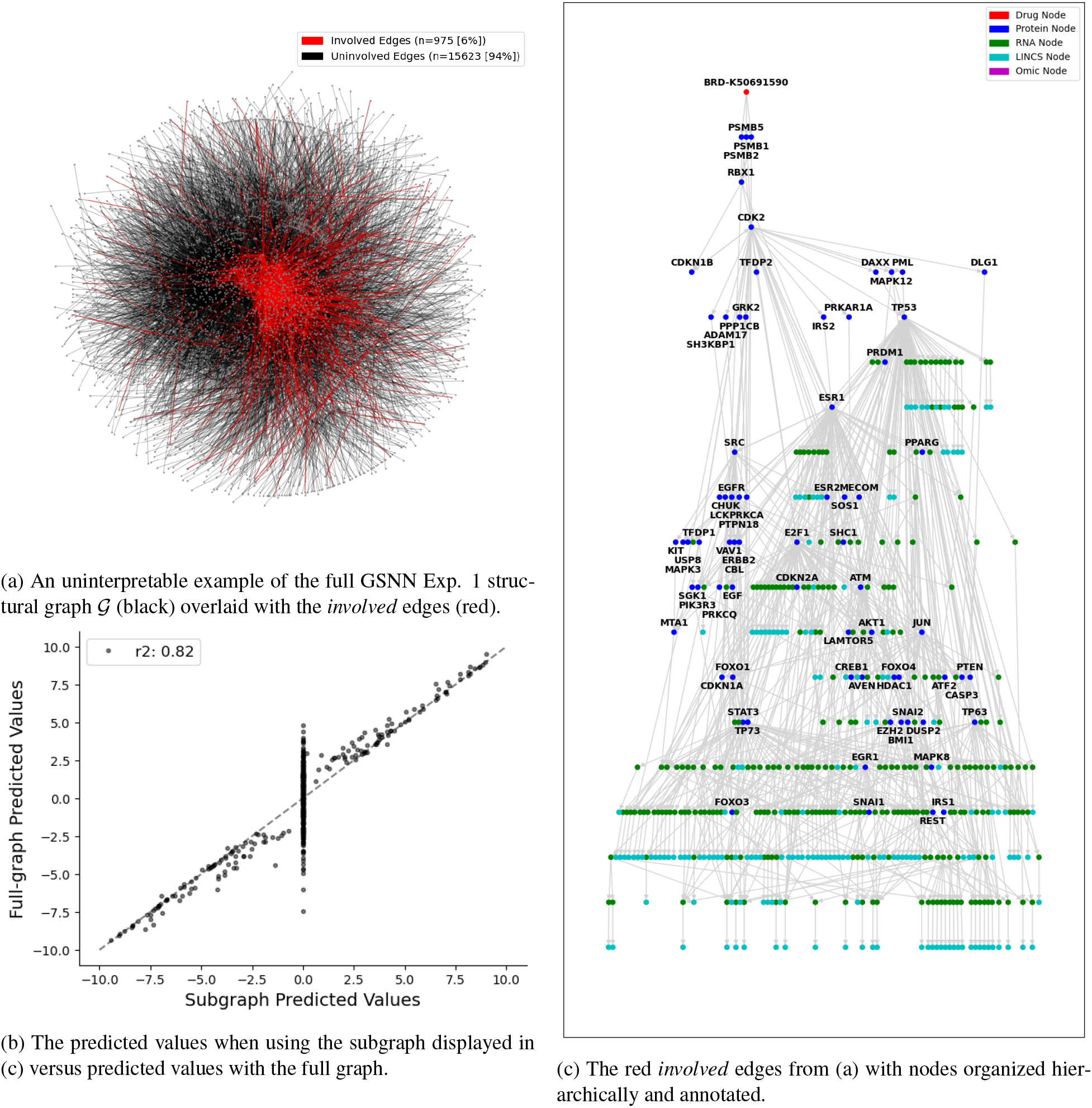
The *GSNNExplainer* method can be used to identify a subgraph that maintains comparable prediction outputs. This can help delineate which edges are involved in the prediction of a given outcome. In this example, we explain the prediction of gene expression response to a proteasome inhibitor (BRD-K50691590) in the PC3 cell line (prostate cancer). For this observation, predictions using the subgraph shown in (b) maintains 82% of the variance of the full-graph prediction.

Since the GSNNExplainer utilizes the stochastic softmax-gumbel distribution to induce discrete edge selection, one concern is the repeatability of the edge importance scores generated by subsequent GSNNExplainer replicates. We investigate this concern by running replicates (n=5) and computing the Pearson correlation of subsequent replicates; the results of a representative use case are shown in Table 5. On average, the pairwise correlation between replicates for this example is greater than 0.9, suggesting repeatable results. The small amount of discordance between replicates is likely due to a tendency for the GSNNExplainer to focus on outputs with larger prediction values, thus ignoring low-value predictions (which are more likely to be dominated by measurement noise). Considering the abundance of predicted outputs, it is likely that there is a stochastic inclusion of low-value predicted outputs. Repeatability between replicates of the GSNNExplainer can be further improved by averaging the edge importance scores between replicates; for example, the pairwise averaging of replicates in Table 5 improves the average pairwise correlation to 0.96. This result suggests that, for applications where repeatability is critical, multiple GSNNExplainer replicates should be computed and the results averaged.

**Table 5:**
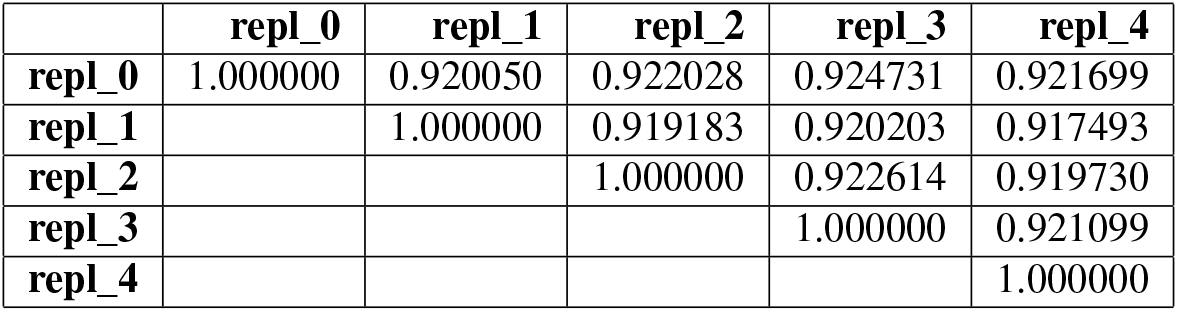
The Pearson correlation between importance scores generated by subsequent GSNNExplainer replicates (n=5). Performed on Dabrafenib (BRD-K09951645) in A375 cell line at a 10uM dose. The average pairwise correlation between replicates is 0.92.

|  | repl_0 | repl_1 | repl_2 | repl_3 | repl_4 |
| --- | --- | --- | --- | --- | --- |
| repl_0 | 1.000000 | 0.920050 | 0.922028 | 0.924731 | 0.921699 |
| repl_1 |  | 1.000000 | 0.919183 | 0.920203 | 0.917493 |
| repl_2 |  |  | 1.000000 | 0.922614 | 0.919730 |
| repl_3 |  |  |  | 1.000000 | 0.921099 |
| repl_4 |  |  |  |  | 1.000000 |

### 5.4 Evaluation of predicted cell viability

Perturbed expression can be useful to characterize the multifaceted response of a biological system; however, it is sometimes convenient to measure a perturbation’s response by a simple phenotypic outcome. For example, the field of *cancer drug response* (CDR) predominantly uses cell viability (or summary metrics) as the outcome variable. Past research has shown that perturbed expression can be used to identify cell death signatures, and that perturbed expression can be used to accurately predict cell viability [87, 57]. In this section, we train a deep ensemble of probabilistic 1-layer neural networks (*f*_*viab*_) to predict cell viability from predicted perturbed expression.

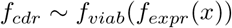

Where *f*_*expr*_ is a frozen^11^ GSNN or NN model from Exp. 1. We optimize *f*_*viab*_ using single-agent cell viability data from the PRISM dataset, and evaluate on a hold-out test dataset (62 cell lines, 305 drugs, ∼115k observations). We then use the NCI ALMANAC dataset [37] to evaluate the performance when applied to unseen two-drug combinations (13 cell lines, 24 drugs, 264 combinations, ∼30k observations). The primary purpose of this evaluation is to compare the relative usefulness of the expression predictions made by the GSNN or NN models. If the GSNN predictions are more robust and mechanistically grounded, then we expect them to perform better in downstream applications such as prediction of cell viability (single and combination agents).

The results of cell viability predictions for single-agent and two-drug combination are shown in Table 6. We find that single-agent cell viability predictions from a GSNN model perform comparably to predictions made with a NN model; however, the GSNN model markedly outperforms the NN model when predicting cell viability for two-drug combinations with a mean squared error (MSE) of less than half the NN model. To evaluate the calibration of cell viability predictions (i.e., the quality of uncertainty quantification), we use the expected calibration error (ECE) [29] and the correlation between predicted variance and error (EVC). In Figure 9 we compare the performance GSNN vs NN predictions when grouped within two-drug combinations (i.e., performance calculated over all doses and cell lines tested with a given drug combination). The GSNN has lower MSE for almost all tested drug combinations with the notable exception of combinations involving the prodrug^12^ Romidepsin, which inhibits histone deacetylases (HDACs) [96].

**Table 6:** Predicted cell viability performance. Values in bold indicate the best performance. The column acronyms are Mean Squared Error (MSE), Expected Calibration Error (ECE), Error-Variance Correlation (EVC), Accuracy (Acc), Area under the receiver operator curve (AUROC). The “Null/Rand.” row refers to metrics computed using random predictions (uniform(0,1)).

| $f_{expr}$ arch. | $f_{viab}$ arch. | MSE | r (pearson) | ECE |
| --- | --- | --- | --- | --- |
| GSNN | NN | **0.017** | 0.87 | 0.11 |
| NN | NN | 0.022 | **0.90** | **0.11** |
| Null/Rand. | Null/Rand. | 0.263 | -0.02 | NA |

(a) Performance evaluated on a hold-out test set of single agent drugs (PRISM dataset).
| $f_{expr}$ arch. | $f_{viab}$ arch. | MSE | r (pearson) | ECE | EVC | Acc. ($y, \hat{y} > 0.5$) | AUROC ($y > 0.5$) |
| --- | --- | --- | --- | --- | --- | --- | --- |
| GSNN | NN | **0.16** | **0.30** | **0.28** | **0.22** | **0.75** | **0.7** |
| NN | NN | 0.34 | 0.22 | 0.35 | -0.24 | 0.46 | 0.63 |
| Null/Rand. | Null/Rand. | 0.27 | 0.00 | NA | NA | 0.5 | 0.5 |

(b) Performance evaluated on unseen two-agent drug combinations (NCI Almanac dataset).

**Figure 9:**
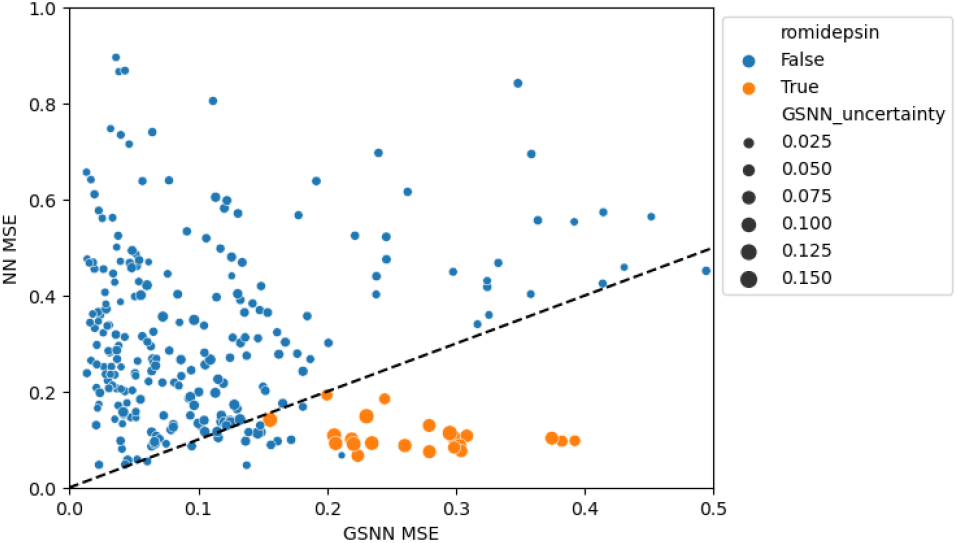
Cell viability prediction error (MSE) computed within drug combinations. X-axis characterizes GSNN error and Y-axis characterizes NN performance; all drug combinations above the diagonal (black dashed line) are better predicted by the GSNN. Notably, all combinations with Romidepsin (orange) were poorly predicted by the GSNN. The size of the points indicates the predicted uncertainty of the GSNN (mean variance computed within combination), which should correlate with GSNN error if the predictions are well calibrated.

It is important to acknowledge that training and evaluating a model on different datasets can introduce several limitations. For example, a seemingly marginal performance of ∼0.3 Pearson correlation should be interpreted in the context of potential data issues. This consideration is particularly relevant to cell viability measurements, which are known to suffer from limited reproducibility and measurement noise [66]. Additionally, the datasets used in our study, namely the single agent data from PRISM [21] and the combination agent data from the NCI Almanac [1, 37], were generated using different cell viability assays (PRISM and NCI-60 protocol, respectively). This disparity raises the possibility of covariate shift [71], which may affect our results. To address this concerns, we refined our evaluation by binarizing cell viability (y > 0.5) and employing classification metrics, specifically accuracy (Acc.) and area under the receiver operator curve (AUROC). The results in Table 6 show that the GSNN model demonstrated superior performance across all classification metrics when predicting cell viability of two-drug combinations.

### 5.5 Disease-specific drug prioritization

In this section, we use the GSNN model (same hyperparameters as Exp. 1) to rank drugs by their selective response in cell lines derived from specific cancer types. We then evaluate the rationality of these prioritizations by comparing them with a limited number of FDA-approved drug indications. We manually select the target and background diseases in Table 7 by choosing diseases that have cell lines in the GSNN cell-space and indications in the PRISM drug repurposing dataset [21]. We use Monte Carlo simulations (N=1000) to estimate the probability that target cell lines have lower average cell viability than background cell lines (*p*_*sens*_). We then rank drugs by *p*_*sens*_ and report the performance of our ranking in Table 7. The reported p-values are calculated as the proportion of null-AUROC values (e.g., random drug lists; N=1000) that are greater than our AUROC value. The GSNN prioritization performs well across disease types with perfect AUROC values in 8/10 disease-specific prioritizations; however, this approach would benefit from a greater number of drug indications to accurately measure significance. For example, Acute Myeloid Leukemia (AML) and Prostate Cancer have only a single drug with indication for these diseases, and prioritizations involving these diseases do not have significant p-values (alpha=0.05) even though they achieved a perfect AUROC score (i.e., all drugs with the target disease indication are prioritized before drugs with the background disease indication). These results should also be interpreted with caution, as the drug indications in the PRISM drug repurposing dataset may not be up-to-date or may not capture ongoing research. For instance, when prioritizing drugs that are preferentially cytotoxic in breast cancer compared to non-small cell lung cancer (NSCLC) our algorithm ranks Afatinib and Gefitinib (EGFR inhibitors with indications for NSCLC) above drugs with breast cancer indications. Although this initially appears to be an inaccurate ranking, a literature search reveals evidence supporting both Afatinib and Gefitinib as a potential treatment option for breast cancer [39, 54, 7]. In prioritization for selective response in NSCLC over Kidney Cancer, Axitinib (with indication for renal cell carcinoma) was ranked above drugs with indication for NSCLC (alectinib, certitinib), but there is research suggesting that axitinib may also be useful for the treatment of NSCLC [78]. Additionally, a systematic review of ALK inhibitors (including ceritinib, alectinib; indications for NSCLC) has shown early promising results for use in renal cancer [40]. It may be that similar diseases will have missing or overlapping disease indications that can confound the evaluation of our drug prioritization.

**Table 7:** The results of disease-specific drug prioritizations evaluated on FDA approved drug indications. Bold font indicates the greatest AUROC value. The last two columns characterize the number of drugs with indications for target or background diseases, respectively. Note that we define the background cell lines as those from a specific disease rather than all non-target cell lines to ensure distinct cellular contexts and clear prioritization goals.

| Target Dis. (# lines) | Background Dis. (# lines) | GSNN AUROC (FDR) | NN AUROC (FDR) | # target indications | # background indications |
| --- | --- | --- | --- | --- | --- |
| NSCLC (8) | AML (2) | **1.00** (0.17) | 0.60 (0.50) | 5 | 1 |
| breast (9) | prostate (3) | 1.00 (0.20) | 1.00 (0.19) | 4 | 1 |
| breast (9) | AML (2) | 1.00 (0.20) | 1.00 (0.20) | 4 | 1 |
| NSCLC (8) | prostate (3) | **1.00 (0.17)** | 0.60 (0.47) | 5 | 1 |
| breast (9) | NSCLC (8) | **0.80 (0.10)** | 0.75 (0.15) | 4 | 5 |
| melanoma (7) | breast (9) | 1.00 (0.02) | 1.00 (0.02) | 4 | 4 |
| breast (9) | kidney (2) | 1.00 (0.07) | 1.00 (0.07) | 4 | 2 |
| melanoma (7) | NSCLC (8) | 1.00 (0.01) | 1.00 (0.01) | 4 | 5 |
| melanoma (7) | kidney (2) | 1.00 (0.07) | 1.00 (0.06) | 4 | 2 |
| NSCLC (8) | kidney (2) | 0.70 (0.28) | **0.90 (0.10)** | 5 | 2 |

## 6 Discussion

As suggested by the NFL theorem and further supported by the success of problem-specific models in vision (CNNs) and NLP (Transformers) tasks, including inductive biases into deep learning algorithms is an effective way to improve performance. In this work, we have shown that *graph structured neural networks* can use prior knowledge, encoded as a graph, to impose useful inductive biases. When applied to a perturbation biology task, the GSNN algorithm outperforms traditional neural networks, graph neural networks (GCN, GAT, and GIN) and GSNN models operating on randomized biological networks. We further investigate *local* performance and show that the GSNN model performs particularly well in a subset of drugs, genes, doses, and diseases. Variation in local performance may suggest that the GSNN algorithm is better suited to model certain types of perturbation, biological processes, or cellular contexts. An in-depth understanding of these advantages or pitfalls could be used to refine the GSNN algorithm or the choice of prior knowledge to build even more robust perturbation models.

The proposed *GSNNExplainer* procedure enables biologically relevant explanation of GSNN predictions, which can be used as a testable hypothesis to build knowledge of cell signaling processes or to develop trust in the GSNN algorithm. Furthermore, the predictions made by GSNNs are both traceable^13^ and inspectable^14^. Future work should also investigate how model-agnostic interpretability methods, such as SHAP [58], can be used to understand the behavior of specific function nodes and could be used to identify the specific role a molecular entity plays in signaling. These aspects are important improvements over traditional deep learning and are a necessary step toward the development of trustworthy models of perturbation biology. Well-validated GSNN models have the potential for application in basic research, pre-clinical drug prioritization, and as a clinical decision aid.

We have shown that the GSNN method can be used to effectively predict disease-specific drug prioritizations, and this suggests that these methods could be used for a wide range of prioritization goals that are uniquely tailored to specific research goals. Furthermore, we have shown that the GSNN can accurately predict the cell viability of drug combinations when trained on single agents and outperforms equivalent models that use traditional NNs. These results suggest that the GSNN algorithm could also be used for prioritization of drug combinations.

### 6.1 Limitations

#### 6.1.1 Appropriate inclusion of molecular entities and interactions

The introduction of prior knowledge in the GSNN can be both a boon and a curse: choosing the right set of molecular entities and interactions is liable to create interpretable high-performing models; but choosing the wrong set and we may over-constrain the model, resulting in poor models and inaccurate prediction logic. Identification of the correct subset of molecular entities required to model a system is a challenging task. In the methods we presented, we used Reactome pathways to select molecular entities that we believe are critical to certain processes and which we can tailor to the type of drug or disease we wish to model. While we believe this is justified as a naive first approach, we do note that there are many ways this approach is lacking. Many knowledgebases, including Reactome, partition healthy signaling pathways separate from diseased signaling pathways, and therefore inclusion of relevant healthy pathways may miss molecular entities critical to disease. In future work, we would like to address this by expert curation or methodological development of entity and interaction selection algorithms. An attractive research direction is to perform hyper-optimization of the biological network during model training. For example, reinforcement learning could be used to select the best set of molecular entities and interactions that maximize the performance of a set of observations.

In this work, we used the *Omnipath* resource for construction of our prior knowledge graph. A limitation that may arise from this is that the Omnipath resource focuses primarily on protein-protein, protein-DNA, and protein-RNA interactions and does not include some relevant molecular entities, such as ions. Ions, such as calcium or potassium, are well known to play an important role in many signaling pathways [28]. Further work may wish to explicitly model ion-like molecular entities; however, due to the role in many different interactions and pathways, encoding them as a distinct entity (i.e., function node) is likely to result in a high centrality^15^ that may lead to inappropriate modeling of pathway cross-talk. For example, two unrelated pathways (e.g., different cellular compartments) that both involve calcium ions would have an unintended means of signaling. This challenge may suggest the need for additional learning mechanisms to model ion-like entities, or any entity that is independently involved in many pathways.

The GSNN method cannot learn de-novo molecular entity interactions and a consequence of this is that understudied pathways are likely to lack appropriate prior knowledge to be appropriate for use with the GSNN. Additionally, the inability to learn new entity relationships makes the GSNN method critically dependent on user selection of accurate interactions and, therefore, susceptible to over-constraining. To address these limitations, future work should consider how to infer new relationships between entities during model training. For example, a simple method would be the incorporation of a “global” node, to which all entities are connected. This addition would enable inference of new relationships between entities *through* the global node. Regularization might be applied to balance exploration of new interactions and exploitation of high-fidelity known interactions.

#### 6.1.2 The drug-target perturbation premise

As described in section 2, we adopt a drug-target premise of chemical perturbation, which requires knowledge and presence of the proteins that a drug binds to; however, some drug mechanisms indirectly affect protein signaling. For instance, some molecular mechanisms change cellular conditions (which we term “condition-response” perturbations), which then activate protein sensors that measure the changing environment and initiate a response; examples of this include hypoxia, heat shock response, oxidative stress, osmotive stress, and DNA damage response. Hypoxia-inducible factors (HIFs) measure oxygen levels and activate transcriptional responses to adapt [95], PARP and DNA-PK proteins initiate the DNA damage response (DDR) signaling pathway [55]. DNA damaging drugs common in cancer therapies, such as Cisplatin, bind directly to DNA and cause DNA adducts [4]. As such, Cisplatin does not have any primary protein targets and yet it is likely to lead to a DDR or apoptotic response. Future work should identify alternative ways to encode these condition-response perturbations. A simple approach may be to include intermittent *process* nodes, for example, include a “DNA damage” node and structure the network so that *Drug* → *DNA damage* → *Protein Signaling Cascade*; however, this approach will require significant expert knowledge and manual curation.

#### 6.1.3 Limitations of the LINCS L1000 dataset

The LINCS L1000 dataset [85] is a large, high-throughput data set that characterizes thousands of drugs in hundreds of cell lines, however, it has known data quality issues [16, 70, 53, 17, 23]. Future work in this field would benefit from the application of the GSNN algorithm to alternative datasets such as the Cancer Perturbed Proteomics Atlas (CPPA) [105]. Ensuring that the GSNN performs well on a range of datasets would provide additional evidence that this algorithm can be applied effectively to model cellular signaling.

#### 6.1.4 Scalability and re-usability

The current formulation of the GSNN algorithm is an order of magnitude slower than traditional neural networks (see supp. 9.3), and therefore it can be computationally expensive to apply to large biological networks or datasets. There are several approaches that could improve GSNN training and inference speeds. In Section 4.9 we describe how the GSNNExplainer can be used to identify a subset of edges that are required to predict an observation. Using analogous methods to prune unimportant edges during the training process or at inference time could significantly reduce the compute requirements. Moreover, each drug perturbation can only affect downstream nodes, and many nodes are likely to be inaccessible to a given drug. This constraint suggests that we could obtain equivalent performance with drug-specific forward passes, which operate on a subset of the full biological graph and would markedly reduce the compute requirements. This concept of drug- or observation-specific forward passes also suggests the premise of *reusuability*. For instance, a function node could be trained using one datatype, network, pathway, or drug set, and then *re-used* in a new datatype or pathway. This approach could enable efficient *localized* training of much larger cellular or microenvironment models.

#### 6.1.5 Appropriate parameter sharing in GSNNs

The GSNN algorithm overcomes many limitations of GNNs by eschewing the parameter-sharing paradigm of GNN message aggregation. While we have shown that our approach has notable advantages for modeling cell signaling, there are also limitations to our approach. Foremost, parameter sharing can significantly reduce the number of trainable weights and, consequently, may improve generalizability in some prediction tasks. This work is likely to benefit from future research that identifies aspects of cellular signaling where parameter sharing may be appropriate and useful, as well as to reinforce aspects where it is not. For instance, the relationship between ‘omic features and molecular state of respective entites is likely to be similar across much of the interactome (e.g., deleterious mutations are likely to prevent the involvement of the respective proteins). Given this consideration, we believe that using parameter sharing to infer the state of molecular entities (i.e., cellular context) could be particularly useful in reducing the number of trainable weights and improving the performance of the GSNN algorithm.

#### 6.1.6 Pathway isolation via protein localization

Cell signaling is typically characterized by biological pathways, which can function independently or interact with each other (pathway *cross-talk*). For a protein-protein interaction (PPI) to be active, the involved proteins must not only be structurally compatible but also be co-localized. Proteins from different pathways may be found in separate sub-cellular compartments, which can prevent interaction and pathway cross-talk. However, interaction between these isolated pathways can occur by movement of a protein between compartments (e.g., translocation of transcription factors to the nucleus).

Our current method for constructing biological networks does not consider the specific pathway or sub-cellular location of proteins. This oversight may lead to inaccuracies in understanding inter-pathway interactions. For instance, if our model merges two pathways that share a protein but are otherwise isolated, it might incorrectly suggest inter-pathway interactions that do not actually occur.

The GSNN algorithm was designed to identify and respond to different input signals (i.e., *source awareness*), suggesting that it could learn specific pathway responses with the appropriate training data. However, given the current constraints of perturbation biology^16^, the GSNN algorithm may improve with additional inductive biases that promote signaling within specific pathways or sub-cellular areas. One way to do this is by using pathway or location annotations to clarify the role and position of each protein. In this approach, a protein could be represented by several function nodes, each modeling the protein’s role in a different pathway or cellular location. Including edges between the same molecular entity in different compartments would allow for pathway cross-talk via translocation (e.g., NFkB_cytosol <-> NFkB_nucleus), and inter- vs intra-compartment signaling could be mediated by regularization (e.g., weight decay) on between-compartment edge weights.

#### 6.1.7 Modeling time with Graph Structured Neural Ordinary Differential Equations

In its current state, the GSNN algorithm only predicts one time point (24H); however, future work should pursue methods to include time series prediction, which better aligns with the true behavior of transcriptional response and would improve model usefulness. Neural Ordinary Differential Equations (ODE) have been proposed to solve ODEs using neural networks [15, 69], however, parameterizing the neural ODE is likely to fall victim to many of the limitations we discussed for traditional neural networks. The GSNN architecture is well suited to parameterize the neural ODE, which would allow the inclusion of prior knowledge constraints and could be applied to time-series modeling of transcriptional response. In practice, the GSNN layers would be replaced by sequential ODE solver steps to compute the change in state over time.

#### 6.1.8 GSNNs for multi-cellular models

Another attractive future direction is the inclusion of multi-cellular or microenvironment models to study more complex behaviors of an organism. In simplicity, two cells could feasibly be encoded as a single GSNN model, with known cell-cell interactions connecting the two distinct cellular models. Such an approach could be used to study the impact of immune cells or tumor microenvironment on drug response. Ignoring the current pragmatic constraints (e.g., memory and compute requirements), the GSNN algorithm may one day be used to model drug response at the tissue or multi-tissue scale; modeling the involvement of hundreds or thousands of cells.

#### 6.1.9 Inclusion of experimental conditions

In-vitro cellular models of drug response require a variety of experimental conditions particular to the assay and cellular model, and these conditions are liable to introduce complexities and bias to the measured response. For example, many drug assays including the LINCS L1000 assay use dimethylsulfoxide (DMSO) to solubilize the various drugs that are tested. There is evidence that DMSO causes statistically significant expression changes in many pathways and cellular models [6]. Furthermore, the Drugbank database lists three known protein targets for DMSO (Accession Number: DB01093), including the MYC transcription factor [96]. To adjust for these experimental conditions, LINCS data processing uses zero-drug DMSO replicates as control when calculating gene perturbations. Even with controls, the various experimental conditions may introduce unexpected biases. Including experimental conditions as inputs to the model could better account for these factors and may help improve model performance and reliability. The presence of DMSO, for instance, could be encoded in the drug-target premise and therefore each observation would be a combination of DMSO + drug. Including experimental conditions in this way may help delineate the expression changes induced by DMSO from the changes induced by the drug.

#### 6.1.10 Encoding differential RNA splicing

In it’s current formulation, we chose to construct a simple biological network where DNA and RNA are modeled as a joint entity. This approach is memory efficient and captures much of the prior knowledge; however, future work may benefit from the development of biological networks where DNA and RNA are modeled separately. Such an approach would allow for the inclusion of multiple RNA transcripts from the same DNA and would enable a more detailed characterization of the molecular landscape. A current limitation of this approach is that most molecular interaction databases are characterized at the gene level, which prevents a detailed understanding of how these interactions change with differential splicing.

## 7 Data and Code Availability

The GSNN project codebase is available in the github repository: https://github.com/nathanieljevans/GSNN. All datasets used in this project are publicly available; for further documentation and precise sources, see the “get_data.sh” file.

## 8 Acknowledgements

This work was supported by the National Cancer Institute (1U54CA224019). N.E. was supported by fellowship funding from the National Cancer Institute (T32 CA106195), National Center for Advancing Translational Sciences (TL1TR002371) and the National Institute of Allergy and Infectious Diseases / National Library of Medicine (NLM) (5T15LM007088).

The authors would like to acknowledge the Oregon Health & Science University department of medical informatics and clinical epidemiology (DMICE) faculty and students of the Oregon Health & Science University Department of Medical Informatics and Clinical Epidemiology (DMICE) who have made this work possible. In particular, we would like to thank Justine Ngyuen, Jennifer Sullivan, Ben Cordier, Chris Klocke, and Gareth Harman for their feedback on the project.

## 9 Supplementary

### 9.1 Experiment Details

Each experiment is described by a set of hyperparameters that specify the nodes (drugs, proteins, RNA & LINCS genes), cell lines, and observations that will be included. One of the key parameters is the choice of proteins to be included in the biological graph on which the GSNN will operate. Ideally, we would select a subset of proteins that are relevant to a certain drug or set of drugs; however, inferring which proteins are involved in a given drug response is a challenging task. In lieu of identifying proteins that are relevant to the drug response, we select proteins based on their involvement with specific biological processes or pathways and manually select the subset of pathways that we believe are likely to be involved in the drug response. In experiments 1-3, we first select a set *primary* Reactome pathways and then manually comb through each primary pathway to identify *linked* pathways^17^. To avoid exceptionally large protein sets, we use our discretion to select *linked* pathways which we believe are relevant to the *primary* pathway. For instance, we generally avoided including pathways that are unlikely to be well modeled by the GSNN premise of cellular signaling such as DNA damage response^18^. We also did not include disease pathways, which Reactome has designated as separate pathways; therefore, all specified pathways should be considered canonical and healthy signaling processes. The experiment pathway details are shown in Table 8. Although each experiment has a unique set of *primary* pathways, there are many included *linked* pathways which are shared by all three experiments. Additionally, many proteins have multiple roles in several pathways. This overlap in pathways and protein roles means that even distinct experiment pathway parameters may result in relatively similar biological networks. Figure 10 shows the overlapping entities between experiments 1-3. Of note, while most elements have substantial overlap between each experiment, the protein-space^19^ is relatively distinct between each experiment.

**Table 8:**
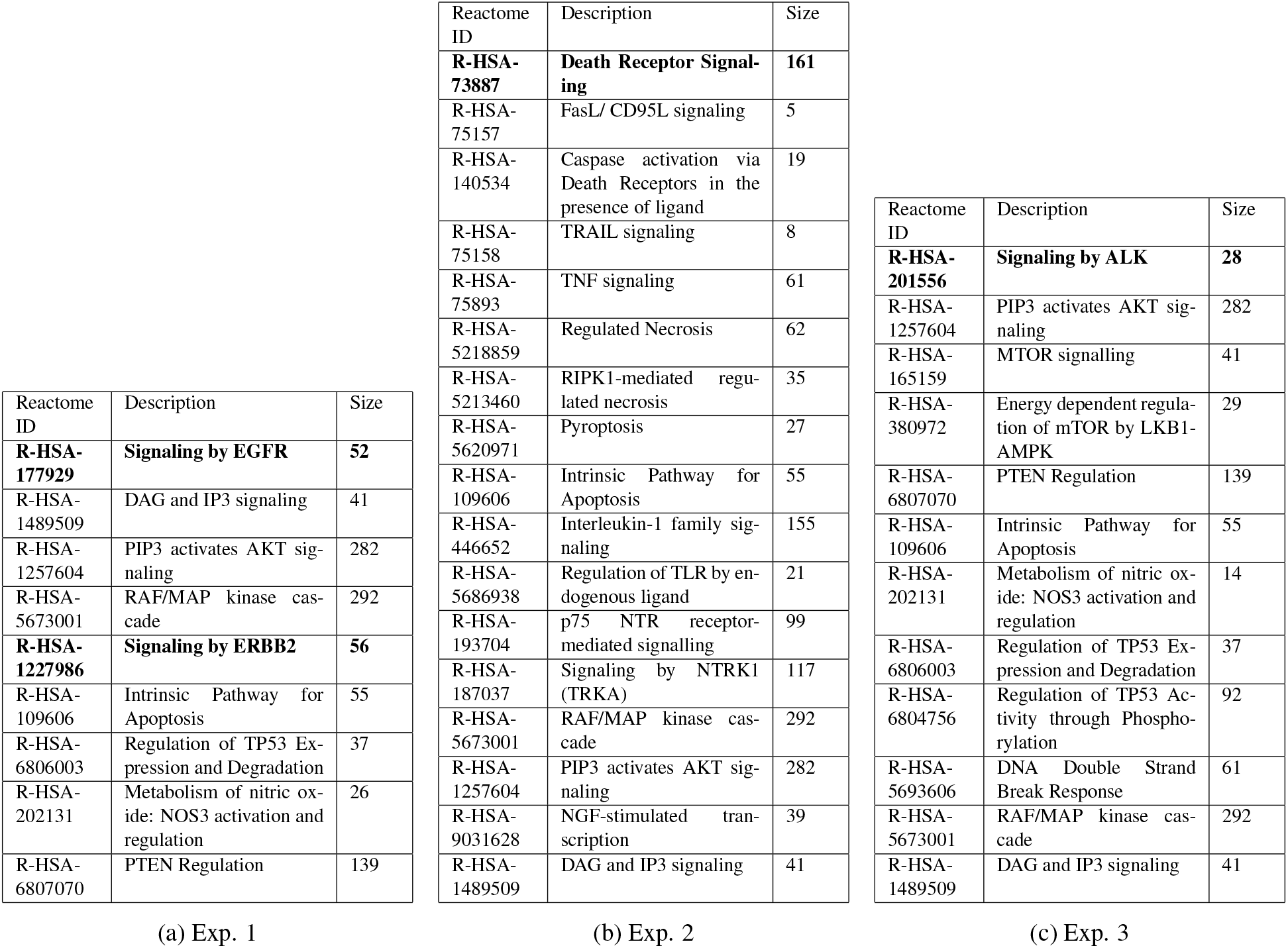
(a-c) The pathways that were used in each experiment to specify the proteins included in the GSNN input graph. Bold text indicates the initial pathway choice from which all other pathways were “linked.” Pathway size refers to the number of proteins in each reactome pathway and may not reflect the exact number of proteins included in the resulting biological network.

**Figure 10:**
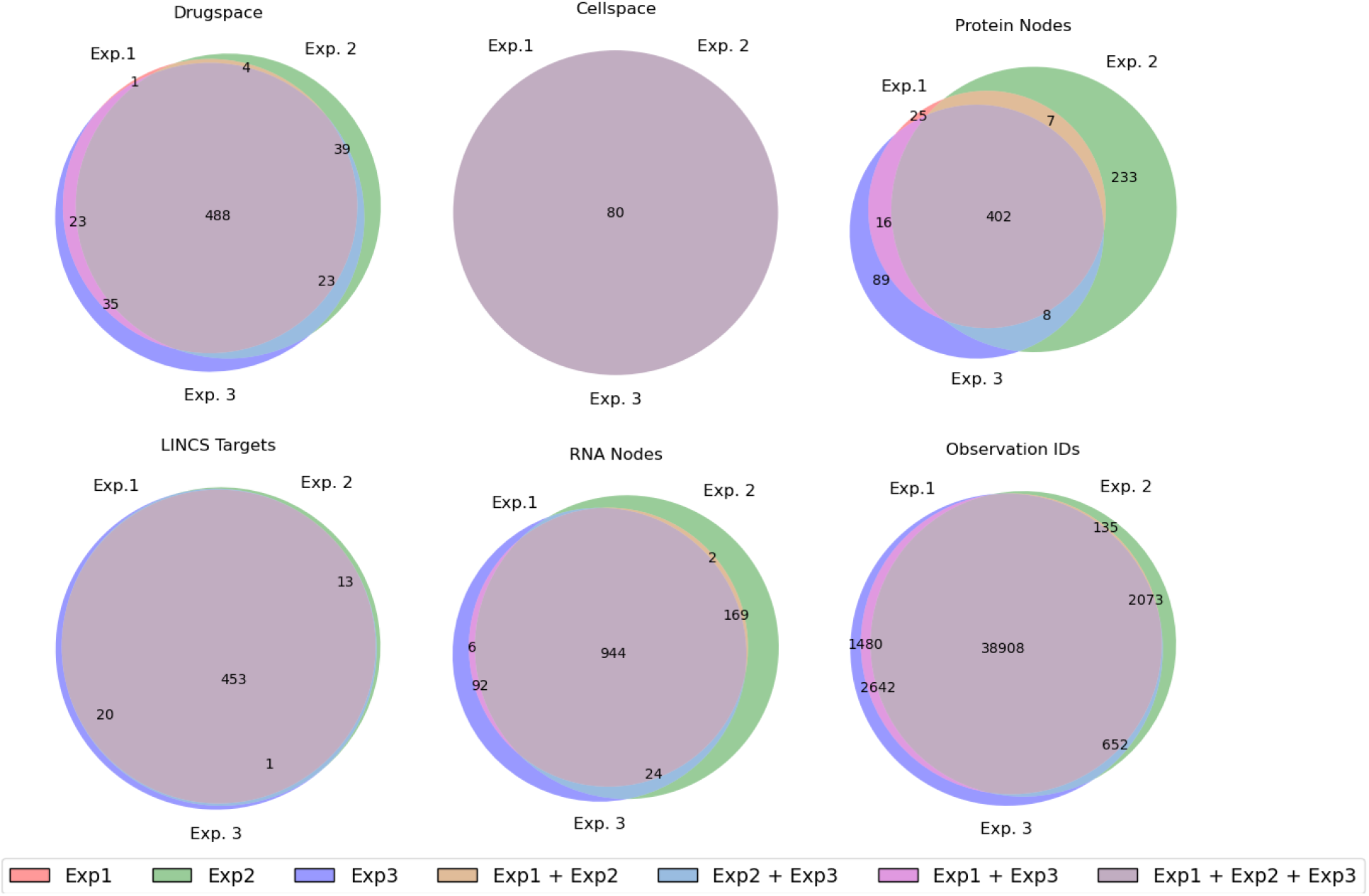
The overlapping elements of each experiment. Most of the targets, nodes and drugs are shared across all three experiments; however, there are distinct protein subsets for each of the three experiments.

### 9.2 Number of model parameters: GSNN vs. NN

In table 9 we report the number of GSNN and NN parameters in the best-performing models of each experiment. Across all three experiments, the best GSNN models from each fold had more trainable parameters than the best NN model from that fold. This may be indicative of the GSNN model being less prone to over-fitting. Another explanation is that, due to the GSNN biological graph structure, there are likely to be many function nodes that are rarely involved in prediction logic or impact only a few targets (i.e., only a few LINCS nodes are descendants) and therefore the functional set of parameters may not be well represented by the total number of trainable parameters. In other words, prior knowledge may lead to some function nodes being effectively spurious or underutilized, and therefore the direct parameter comparison should be interpreted with caution.

**Table 9:** Number of trainable parameters of the GSNN and NN algorithms used in experiments 1-3 (median of best models from each MCCV fold). Percent change is calculated as 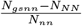, where *N* is the median number of algorithm parameters

| EXP. ID | Num. GSNN params | Num. NN params | Percent Change |
| --- | --- | --- | --- |
| exp1 | 6.56e+06 | 2.23e+06 | 193.8 % |
| exp2 | 7.85e+06 | 5.68e+06 | 38.2 % |
| exp3 | 6.70e+06 | 5.3e+06 | 26.5 % |
| AVG. |  |  | 86.2 % |

### 9.3 Computational Complexity of the GSNN method

The GSNN algorithm takes significantly longer to train due to being a particularly deep architecture and due to it’s use of sparse matrix operations. Table 10 reports the average training times for each algorithm. Specifically, the GSNN algorithm requires between 3-15 times as much training time as the alternative algorithms tested (NN, GNN). Of note, however, are the training curves shown in Figure 11 that compare the validation performance by epoch for representative GSNN and NN models; The GSNN validation performance increases markedly faster, achieving approximately the maximum NN performance in the first 20 epochs. This aspect of the training dynamics may suggest that the GSNN algorithm can be trained with fewer epochs, which would markedly reduce the compute requirements.

**Table 10:** Average training time of each algorithm (reported in minutes). Note: GSNN and GNN were trained on GPUs whereas the NNs were trained on CPU only.

| EXP. | GSNN | GNN | NN | GSNN / NN | GSNN / GNN |
| --- | --- | --- | --- | --- | --- |
| exp1 | 419.8 | 105.0 | 29.0 | 14.5 | 4.0 |
| exp2 | 493.3 | 138.4 | 32.4 | 15.2 | 3.6 |
| exp3 | 460.8 | 133.9 | 35.9 | 12.9 | 3.4 |

**Figure 11:**
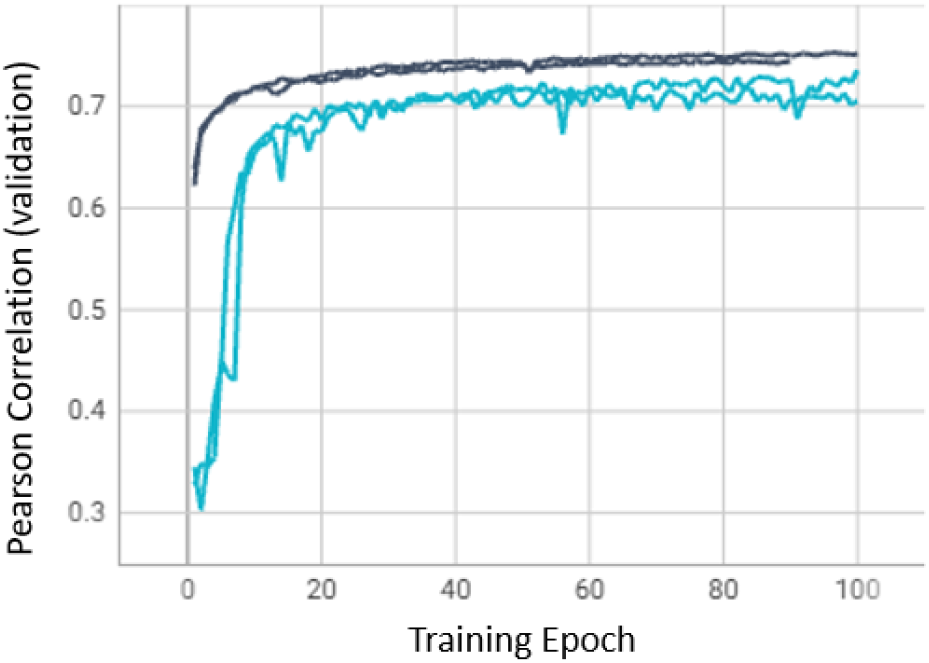
Representative training curves from experiment 1 (EGFR + ERBB2 signaling). Dark Gray/Blue indicates the GSNN training curves, light blue are NN training curves.

### 9.4 Effect of Layer Depth on GSNN performance

The GSNN algorithm passes information during sequential *layers* allowing information to diffuse through the network up to the number of layers L in the model. Cell signaling often involves many entities in many sequential interactions as well as feedback loops that may alter behavior. Due to this trait, deeper networks may be more representative of the underlying biology and therefore more accurate. To test this, we compare the performance of GSNN algorithms with different number of layers (L=10,20). Figure 12 shows the results and suggests that 20-layer GSNNs have a small improvement in performance compared to 10-layer GSNNs. Notably, training deeper neural networks also introduces more parameters, greater memory complexity, and longer training times. It is critical, therefore, that the choice of GSNN layers be tailored to the available hardware and training budget. Improvements in time complexity of the GSNN algorithm may enable deeper and more accurate models.

**Figure 12:**
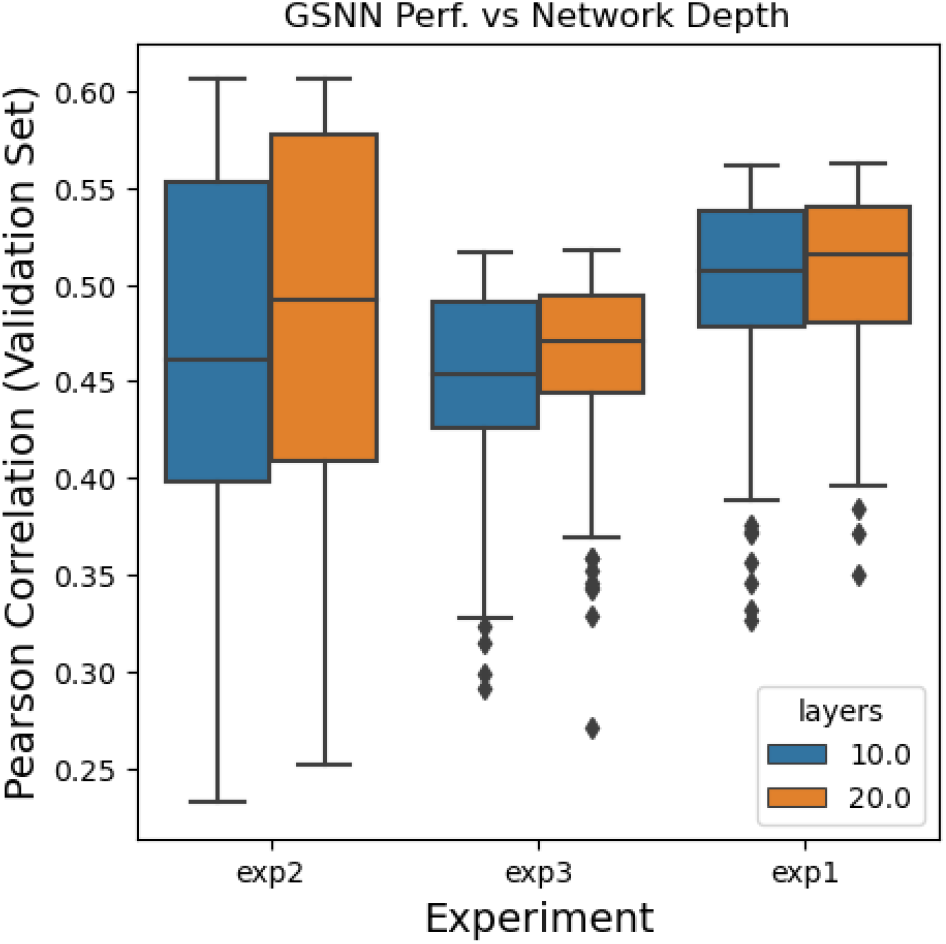
The performance of GSNN algorithms in experiments 1-3 compared by the number of layers (L) hyper-parameter.

## Footnotes

1 The realm of all possible functions

2 An accurate or correct function

3 Referred to as a kernel

4 Note: this gives rise to the assumption of *locality*

5 PAthway Recognition Algorithm using Data Integration on Genomic Models

6 The name *Graph* ***Structured*** *Neural Networks* is intended to recognize the role that SEMs played in the motivation of this work.

7 To avoid ambiguity: there are two “layer” parameters we reference. The function node number of layers (*k* ∼ 1, 2), which describe the number of layers in a function node neural network, and the GSNN’s number of layers (*L* ∼ [10 − 20]) that represent the number of sequential masked linear layers that are used in a GSNN model.

8 Layer here refers to the number of edge-updates, not the number of layers in a function node

9 or metrics derived from cell viability measured over a range of concentrations such as the area under the dose response curve (AUC) or the half maximal inhibitory concentration (IC50)

10 The Benjamini-Hochberg method is used for local performance groupings by cell line, gene, drug and disease where the groups are disjoint sets and therefore the p-values are independent. The Benjamini-Yekutieli method is used for local performance groups by drug-target where an observation may be assigned to two or more groups and therefore p-values are not independent

11 *Frozen* indicates that the parameters of *f*_*expr*_ are not updated during optimization of *f*_*viab*_

12 a drug that is activated once inside the cell.

13 *Traceable* refers to the ability to trace a prediction to intermediary entities, states or input features.

14 *Inspectable* refers to the ability to inspect the behavior of specific intermediate entities

15 centrality is a network science metric characterizing the importance or connectedness to other nodes

16 Perturbation biology data is limited in volume and often measured via noisy assays.

17 Pathways that are not subpathways but are referenced within a pathway, e.g., pathway A -> activates -> pathway B but pathway B is not a subpathway of A)

18 We are concerned that DNA damage drugs will not be well represented by the drug-target premise

19 all proteins included in the biological network

## References

[1] NCI-60 Screening Methodology | NCI-60 Human Tumor Cell Lines Screen | Discovery & Development Services | Developmental Therapeutics Program (DTP).

[2] Trustworthy and responsible AI. Last Modified: 2023-05-04T13:20-04:00.

[3] Arohan Ajit, Koustav Acharya, and Abhishek Samanta. A review of convolutional neural networks. 2020 International Conference on Emerging Trends in Information Technology and Engineering (ic-ETITE), pages 1–5, 2020.

[4] Sara A Aldossary. Review on pharmacology of cisplatin: clinical use, toxicity and mechanism of resistance of cisplatin. Biomedical and Pharmacology Journal, 12(1):7–15, 2019.

[5] Jimmy Lei Ba, Jamie Ryan Kiros, and Geoffrey E. Hinton. Layer normalization, 2016.

[6] Elisa Baldelli, Mahalakshmi Subramanian, Abduljalil M Alsubaie, Guy Oldaker, Maria Emelianenko, Emna El Gazzah, Sara Baglivo, Kimberley A Hodge, Fortunato Bianconi, Vienna Ludovini, et al. Heterogeneous off-target effects of ultra-low dose dimethyl sulfoxide (dmso) on targetable signaling events in lung cancer in vitro models. International Journal of Molecular Sciences, 22(6):2819, 2021.

[7] José Baselga, Joan Albanell, Amparo Ruiz, Ana Lluch, Pere Gascón, Vicente Guillém, Sonia González, Silvia Sauleda, Irene Marimón, Josep M Tabernero, et al. Phase ii and tumor pharmacodynamic study of gefitinib in patients with advanced breast cancer. J Clin Oncol, 23(23):5323–5333, 2005.

[8] Peter W. Battaglia, Jessica B. Hamrick, Victor Bapst, Alvaro Sanchez-Gonzalez, Vinicius Zambaldi, Mateusz Malinowski, Andrea Tacchetti, David Raposo, Adam Santoro, Ryan Faulkner, Caglar Gulcehre, Francis Song, Andrew Ballard, Justin Gilmer, George Dahl, Ashish Vaswani, Kelsey Allen, Charles Nash, Victoria Langston, Chris Dyer, Nicolas Heess, Daan Wierstra, Pushmeet Kohli, Matt Botvinick, Oriol Vinyals, Yujia Li, and Razvan Pascanu. Relational inductive biases, deep learning, and graph networks, 2018.

[9] Yoav Benjamini and Yosef Hochberg. Controlling the false discovery rate: a practical and powerful approach to multiple testing. Journal of the royal statistical society series b-methodological, 57:289–300, 1995.

[10] Yoav Benjamini and Daniel Yekutieli. The control of the false discovery rate in multiple testing under dependency. Annals of Statistics, 29:1165–1188, 2001.

[11] Tanya N Beran and Claudio Violato. Structural equation modeling in medical research: a primer. BMC research notes, 3(1):1–10, 2010.

[12] Aurora S. Blucher, Gabrielle Choonoo, Molly F. Kulesz-Martin, Guanming Wu, and Shannon K. McWeeney. Evidence-based precision oncology with the cancer targetome. Trends in pharmacological sciences, 38 12:1085–1099, 2017.

[13] Aurora S Blucher, Shannon K McWeeney, Lincoln Stein, and Guanming Wu. Visualization of drug target interactions in the contexts of pathways and networks with reactomefiviz. F1000Research, 8, 2019.

[14] Deli Chen, Yankai Lin, Wei Li, Peng Li, Jie Zhou, and Xu Sun. Measuring and relieving the over-smoothing problem for graph neural networks from the topological view, 2019.

[15] Ricky T. Q. Chen, Yulia Rubanova, Jesse Bettencourt, and David Duvenaud. Neural ordinary differential equations, 2019.

[16] L Cheng and L Li. Systematic quality control analysis of LINCS data. 5(11):588–598.

[17] Neil R. Clark, Kevin S. Hu, Axel S. Feldmann, Yan Kou, Edward Y. Chen, Qiaonan Duan, and Avi Ma’ayan. The characteristic direction: a geometrical approach to identify differentially expressed genes. BMC Bioinformatics, 15, 2014.

[18] Djork-Arné Clevert, Thomas Unterthiner, and Sepp Hochreiter. Fast and accurate deep network learning by exponential linear units (elus). arXiv: Learning, 2015.

[19] Cristian Coarfa, Sandra L Grimm, Kimal Rajapakshe, Dimuthu Perera, Hsin-Yi Lu, Xuan Wang, Kurt R Christensen, Qianxing Mo, Dean P Edwards, and Shixia Huang. Reverse-phase protein array: Technology, application, data processing, and integration. Journal of biomolecular techniques: JBT, 32(1):15, 2021.

[20] Steven M. Corsello, Joshua A. Bittker, Zihan Liu, Joshua Gould, Patrick McCarren, Jodi E. Hirschman, Stephen E. Johnston, Anita Vrcic, Bang Wong, Mariya Khan, Jacob K. Asiedu, Rajiv Narayan, C. C. Mader, Aravind Subramanian, and Todd R. Golub. The drug repurposing hub: a next-generation drug library and information resource. Nature Medicine, 23:405–408, 2017.

[21] Steven M Corsello, Rohith T Nagari, Ryan D Spangler, Jordan Rossen, Mustafa Kocak, Jordan G Bryan, Ranad Humeidi, David Peck, Xiaoyun Wu, Andrew A Tang, et al. Discovering the anticancer potential of non-oncology drugs by systematic viability profiling. Nature cancer, 1(2):235–248, 2020.

[22] Lifang Deng, Miao Yang, and Katerina M Marcoulides. Structural equation modeling with many variables: A systematic review of issues and developments. Frontiers in psychology, 9:580, 2018.

[23] Qiaonan Duan, St Patrick Reid, Neil R. Clark, Zichen Wang, Nicolas F. Fernandez, Andrew D. Rouillard, Ben Readhead, Sarah R. Tritsch, Rachel Hodos, Marc Hafner, Mario Niepel, Peter K. Sorger, Joel T. Dudley, Sina Bavari, Rekha G. Panchal, and Avi Ma’ayan. L1000cds2: LINCS l1000 characteristic direction signatures search engine. 2(1):1–12. Number: 1 Publisher: Nature Publishing Group.

[24] Matthias Fey and Jan Eric Lenssen. Fast graph representation learning with pytorch geometric, 2019.

[25] David Freedman, Robert Pisani, and Roger Purves. Statistics (international student edition). Pisani, R. Purves, 4th edn. WW Norton & Company, New York, 2007.

[26] Fabian Fröhlich, Thomas Kessler, Daniel Weindl, Alexey Shadrin, Leonard Schmiester, Hendrik Hache, Artur Muradyan, Moritz Schütte, Ji-Hyun Lim, Matthias Heinig, et al. Efficient parameter estimation enables the prediction of drug response using a mechanistic pan-cancer pathway model. Cell systems, 7(6):567–579, 2018.

[27] Luz Garcia-Alonso, Mahmoud M. Ibrahim, Denes Turei, and Julio Sáez-Rodríguez. Benchmark and integration of resources for the estimation of human transcription factor activities. Genome Research, 29:1363 –1375, 2018.

[28] Marc Gillespie, Bijay Jassal, Ralf Stephan, Marija Milacic, Karen Rothfels, Andrea Senff-Ribeiro, Johannes Griss, Cristoffer Sevilla, Lisa Matthews, Chuqiao Gong, Chuan Deng, Thawfeek Varusai, Eliot Ragueneau, Yusra Haider, Bruce May, Veronica Shamovsky, Joel Weiser, Timothy Brunson, Nasim Sanati, Liam Beckman, Xiang Shao, Antonio Fabregat, Konstantinos Sidiropoulos, Julieth Murillo, Guilherme Viteri, Justin Cook, Solomon Shorser, Gary Bader, Emek Demir, Chris Sander, Robin Haw, Guanming Wu, Lincoln Stein, Henning Hermjakob, and Peter D’Eustachio. The reactome pathway knowledgebase 2022. Nucleic Acids Research, 50(D1):D687–D692, 11 2021.

[29] Chuan Guo, Geoff Pleiss, Yu Sun, and Kilian Q. Weinberger. On calibration of modern neural networks, 2017.

[30] Alexander Sebastian Hauser, Sreenivas Chavali, Ikuo Masuho, Leonie Johanna Jahn, Kirill A. Martemyanov, David E. Gloriam, and M. Madan Babu. Pharmacogenomics of gpcr drug targets. Cell, 172:41–54.e19, 2018.

[31] Winston Haynes. Bonferroni correction. Encyclopedia of systems biology, pages 154–154, 2013.

[32] Kaiming He, Xiangyu Zhang, Shaoqing Ren, and Jian Sun. Deep residual learning for image recognition, 2015.

[33] Kaiming He, Xiangyu Zhang, Shaoqing Ren, and Jian Sun. Delving deep into rectifiers: Surpassing human-level performance on imagenet classification, 2015.

[34] Éléa Héberlé and Anaïs Flore Bardet. Sensitivity of transcription factors to dna methylation. Essays in biochemistry, 63(6):727–741, 2019.

[35] Reinhart Heinrich, Benjamin G Neel, and Tom A Rapoport. Mathematical models of protein kinase signal transduction. Molecular cell, 9(5):957–970, 2002.

[36] Geoffrey Hinton, Oriol Vinyals, and Jeff Dean. Distilling the knowledge in a neural network, 2015.

[37] Susan L Holbeck, Richard Camalier, James A Crowell, Jeevan Prasaad Govindharajulu, Melinda Hollingshead, Lawrence W Anderson, Eric Polley, Larry Rubinstein, Apurva Srivastava, Deborah Wilsker, et al. The national cancer institute almanac: a comprehensive screening resource for the detection of anticancer drug pairs with enhanced therapeutic activity. Cancer research, 77(13):3564–3576, 2017.

[38] Kurt Hornik, Maxwell Stinchcombe, and Halbert White. Multilayer feedforward networks are universal approximators. Neural Networks, 2(5):359–366, 1989.

[39] Sara A Hurvitz, Rebecca Shatsky, and Nadia Harbeck. Afatinib in the treatment of breast cancer. Expert opinion on investigational drugs, 23(7):1039–1047, 2014.

[40] Giovanni Maria Iannantuono, Silvia Riondino, Stefano Sganga, Mario Roselli, and Francesco Torino. Activity of alk inhibitors in renal cancer with alk alterations: A systematic review. International Journal of Molecular Sciences, 23(7):3995, 2022.

[41] Sergey Ioffe and Christian Szegedy. Batch normalization: Accelerating deep network training by reducing internal covariate shift. In International Conference on Machine Learning, 2015.

[42] Eric Jang, Shixiang Gu, and Ben Poole. Categorical reparameterization with gumbel-softmax, 2017.

[43] Ritsert C Jansen. Studying complex biological systems using multifactorial perturbation. Nature Reviews Genetics, 4(2):145–151, 2003.

[44] Kelly Karl, Michael D Paul, Elena B Pasquale, and Kalina Hristova. Ligand bias in receptor tyrosine kinase signaling. Journal of Biological Chemistry, 295(52):18494–18507, 2020.

[45] Thomas Kipf and Max Welling. Semi-supervised classification with graph convolutional networks. ArXiv, abs/1609.02907, 2016.

[46] Brent M Kuenzi, Jisoo Park, Samson H Fong, Kyle S Sanchez, John Lee, Jason F Kreisberg, Jianzhu Ma, and Trey Ideker. Predicting drug response and synergy using a deep learning model of human cancer cells. Cancer cell, 38(5):672–684, 2020.

[47] Michael Kuhn, Christian von Mering, Monica Campillos, Lars Juhl Jensen, and Peer Bork. Stitch: interaction networks of chemicals and proteins. Nucleic Acids Research, 36:D684 – D688, 2007.

[48] Kimberly R Kukurba and Stephen B Montgomery. Rna sequencing and analysis. Cold Spring Harbor Protocols, 2015(11):pdb–top084970, 2015.

[49] Siddharth Krishna Kumar. On weight initialization in deep neural networks, 2017.

[50] Balaji Lakshminarayanan, Alexander Pritzel, and Charles Blundell. Simple and scalable predictive uncertainty estimation using deep ensembles, 2017.

[51] Justin Lamb, Emily D Crawford, David Peck, Joshua W Modell, Irene C Blat, Matthew J Wrobel, Jim Lerner, Jean-Philippe Brunet, Aravind Subramanian, Kenneth N Ross, et al. The connectivity map: using gene-expression signatures to connect small molecules, genes, and disease. science, 313(5795):1929–1935, 2006.

[52] Samuel A. Lambert, Arttu Jolma, Laura F. Campitelli, Pratyush Kumar Das, Yimeng Yin, Mihai Albu, Xiaoting Chen, Jussi Taipale, Timothy R. Hughes, and Matthew T. Weirauch. The human transcription factors. Cell, 172:650–665, 2018.

[53] Zhaoyang Li, Jin Li, and YU Peng. l1kdeconv: an r package for peak calling analysis with lincs l1000 data. BMC Bioinformatics, 18, 2017.

[54] Nancy U Lin, Eric P Winer, Duncan Wheatley, Lisa A Carey, Stephen Houston, David Mendelson, Pamela Munster, Laurie Frakes, Steve Kelly, Agustin A Garcia, et al. A phase ii study of afatinib (bibw 2992), an irreversible erbb family blocker, in patients with her2-positive metastatic breast cancer progressing after trastuzumab. Breast cancer research and treatment, 133:1057–1065, 2012.

[55] Christopher J Lord and Alan Ashworth. The dna damage response and cancer therapy. Nature, 481(7381):287–294, 2012.

[56] Pavel Loskot, Komlan Atitey, and Lyudmila Mihaylova. Comprehensive review of models and methods for inferences in bio-chemical reaction networks. Frontiers in genetics, page 549, 2019.

[57] Jiaxing Lu, Ming Chen, and Yufang Qin. Drug-induced cell viability prediction from lincs-l1000 through wrfen-xgboost algorithm. BMC bioinformatics, 22:1–18, 2021.

[58] Scott M Lundberg and Su-In Lee. A unified approach to interpreting model predictions. Advances in neural information processing systems, 30, 2017.

[59] Jianzhu Ma, Michael Ku Yu, Samson Fong, Keiichiro Ono, Eric Sage, Barry Demchak, Roded Sharan, and Trey Ideker. Using deep learning to model the hierarchical structure and function of a cell. Nature methods, 15(4):290–298, 2018.

[60] Yao Ma, Xiaorui Liu, Neil Shah, and Jiliang Tang. Is homophily a necessity for graph neural networks?, 2023.

[61] Chris J. Maddison, Andriy Mnih, and Yee Whye Teh. The concrete distribution: A continuous relaxation of discrete random variables, 2017.

[62] Warren S McCulloch and Walter Pitts. A logical calculus of the ideas immanent in nervous activity. The bulletin of mathematical biophysics, 5(4):115–133, 1943.

[63] John G Moffat, Fabien Vincent, Jonathan A Lee, Jörg Eder, and Marco Prunotto. Opportunities and challenges in phenotypic drug discovery: an industry perspective. Nature reviews Drug discovery, 16(8):531–543, 2017.

[64] Evan J Molinelli, Anil Korkut, Weiqing Wang, Martin L Miller, Nicholas P Gauthier, Xiaohong Jing, Poorvi Kaushik, Qin He, Gordon Mills, David B Solit, et al. Perturbation biology: inferring signaling networks in cellular systems. PLoS computational biology, 9(12):e1003290, 2013.

[65] Alhassan Mumuni and Fuseini Mumuni. Data augmentation: A comprehensive survey of modern approaches. Array, page 100258, 2022.

[66] Mario Niepel, Marc Hafner, Caitlin E Mills, Kartik Subramanian, Elizabeth H Williams, Mirra Chung, Benjamin Gaudio, Anne Marie Barrette, Alan D Stern, Bin Hu, et al. A multi-center study on the reproducibility of drug-response assays in mammalian cell lines. Cell systems, 9(1):35–48, 2019.

[67] Peter J Park. Chip–seq: advantages and challenges of a maturing technology. Nature reviews genetics, 10(10):669–680, 2009.

[68] Adam Paszke, Sam Gross, Francisco Massa, Adam Lerer, James Bradbury, Gregory Chanan, Trevor Killeen, Zeming Lin, Natalia Gimelshein, Luca Antiga, Alban Desmaison, Andreas Köpf, Edward Yang, Zach DeVito, Martin Raison, Alykhan Tejani, Sasank Chilamkurthy, Benoit Steiner, Lu Fang, Junjie Bai, and Soumith Chintala. Pytorch: An imperative style, high-performance deep learning library, 2019.

[69] Michael Poli, Stefano Massaroli, Junyoung Park, Atsushi Yamashita, Hajime Asama, and Jinkyoo Park. Graph neural ordinary differential equations, 2021.

[70] Yue Qiu, Tianhuan Lu, Hansaim Lim, and Lei Xie. A Bayesian approach to accurate and robust signature detection on LINCS L1000 data. Bioinformatics, 36(9):2787–2795, 01 2020.

[71] Joaquin Quinonero-Candela, Masashi Sugiyama, Anton Schwaighofer, and Neil D. Lawrence. Dataset shift in machine learning. 2022.

[72] Maziar Raissi, Paris Perdikaris, and George E Karniadakis. Physics-informed neural networks: A deep learning framework for solving forward and inverse problems involving nonlinear partial differential equations. Journal of Computational physics, 378:686–707, 2019.

[73] Marco Tulio Ribeiro, Sameer Singh, and Carlos Guestrin. “why should i trust you?”: Explaining the predictions of any classifier, 2016.

[74] Christian P. Robert and George Casella. Monte carlo statistical methods. Technometrics, 47:243 –243, 2005.

[75] Nidhi Sahni, S. Stephen Yi, Mikko Taipale, Juan Fuxman Bass, Jasmin Coulombe-Huntington, Fan Yang, Jian Peng, Jochen Weile, Georgios Ioannis Karras, Yang Wang, István A. Kovács, Atanas Kamburov, Irina Krykbaeva, Mandy Hiu Yi Lam, George Tucker, Vikram Khurana, Amitabh Sharma, Yang-Yu Liu, Nozomu Yachie, Quan Zhong, Yun Shen, Alexandre Palagi, Adriana San-Miguel, Changyu Fan, Dawit Balcha, Amélie Dricot, Daniel M. Jordan, Jennifer M. Walsh, Akash A. Shah, Xinping Yang, Ani K Stoyanova, Alexander T. Leighton, Michael A. Calderwood, Yves Jacob, Michael E. Cusick, Kourosh Salehi-Ashtiani, Luke Whitesell, Shamil R. Sunyaev, Bonnie Berger, Albert-László Barabási, Benoît Charloteaux, David E. Hill, Tong Hao, Frederick P. Roth, Yu Xia, Albertha J. M. Walhout, Susan Lindquist, and Marc Vidal. Widespread macromolecular interaction perturbations in human genetic disorders. Cell, 161:647–660, 2015.

[76] Terence Sanger and Pallavi N. Baljekar. The perceptron: a probabilistic model for information storage and organization in the brain. Psychological review, 65 6:386–408, 1958.

[77] Murat Hüsnü Sazli. A brief review of feed-forward neural networks. 2006.

[78] JH Schiller, J von Pawel, T Larson, SI Ou, SA Limentani, AB Sandler, EE Vokes, S Kim, KF Liau, and PW Bycott. Efficacy and safety of single-agent axitinib (ag-013736; ag) in patients with advanced non–small-cell lung cancer (nsclc): A phase ii trial. Clinical Lung Cancer, 8(7):452, 2007.

[79] Ilana Schlam and Sandra M Swain. Her2-positive breast cancer and tyrosine kinase inhibitors: the time is now. NPJ breast cancer, 7(1):56, 2021.

[80] Gideon Schreiber, Gilad Haran, and Huan-Xiang Zhou. Fundamental aspects of protein-protein association kinetics. Chemical reviews, 109:839–60, 03 2009.

[81] Maya Shamir, Y. Bar-On, Rob Phillips, and Ron Milo. Snapshot: Timescales in cell biology. Cell, 164:1302–1302.e1, 2016.

[82] Jeffrey S Smith, Robert J Lefkowitz, and Sudarshan Rajagopal. Biased signalling: from simple switches to allosteric microprocessors. Nature reviews Drug discovery, 17(4):243–260, 2018.

[83] Eduardo D Sontag and Madalena Chaves. Exact computation of amplification for a class of nonlinear systems arising from cellular signaling pathways. Automatica, 42(11):1987–1992, 2006.

[84] Nitish Srivastava, Geoffrey Hinton, Alex Krizhevsky, Ilya Sutskever, and Ruslan Salakhutdinov. Dropout: a simple way to prevent neural networks from overfitting. The journal of machine learning research, 15(1):1929–1958, 2014.

[85] Aravind Subramanian, Rajiv Narayan, Steven M. Corsello, David Peck, Ted E. Natoli, Xiaodong Lu, Joshua Gould, John F. Davis, Andrew A. Tubelli, Jacob K. Asiedu, David L. Lahr, Jodi E. Hirschman, Zihan Liu, Melanie K. Donahue, Bina Julian, Mariya Khan, David Wadden, Ian Smith, Daniel Lam, Arthur Liberzon, Courtney Toder, Mukta Bagul, Marek Orzechowski, Oana M. Enache, Federica Piccioni, Sarah Johnson, Nicholas J. Lyons, Alice H. Berger, Alykhan F. Shamji, Angela N. Brooks, Anita Vrcic, Corey Flynn, Jacqueline Rosains, David Y. Takeda, Roger Hu, Desiree Davison, Justin Lamb, Kristin G. Ardlie, Larson J Hogstrom, Peyton Greenside, Nathanael S. Gray, Paul A. Clemons, Serena J. Silver, Xiaoyun Wu, Wen-Ning Zhao, Willis Read-Button, Xiaohua Wu, Stephen J. Haggarty, Lucienne V. Ronco, Jesse S. Boehm, Stuart L. Schreiber, John G. Doench, Joshua A. Bittker, David E. Root, Bang Wong, and Todd R. Golub. A next generation connectivity map: L1000 platform and the first 1,000,000 profiles. Cell, 171:1437–1452.e17, 2017.

[86] Mukund Sundararajan, Ankur Taly, and Qiqi Yan. Axiomatic attribution for deep networks, 2017.

[87] Bence Szalai, Vigneshwari Subramanian, Christian H Holland, Róbert Alföldi, László G Puskás, and Julio Saez-Rodriguez. Signatures of cell death and proliferation in perturbation transcriptomics data—from confounding factor to effective prediction. Nucleic Acids Research, 47(19):10010–10026, 2019.

[88] Damian Szklarczyk, Annika L. Gable, Katerina C. Nastou, David Lyon, Rebecca Kirsch, Sampo Pyysalo, Nadezhda T. Doncheva, Marc Legeay, Tao Fang, Peer Bork, Lars Juhl Jensen, and Christian von Mering. The string database in 2021: customizable protein–protein networks, and functional characterization of user-uploaded gene/measurement sets. Nucleic Acids Research, 49:D605 – D612, 2020.

[89] Pauline Traynard, Luis Tobalina, Federica Eduati, Laurence Calzone, and Julio Saez-Rodriguez. Logic modeling in quantitative systems pharmacology. CPT: pharmacometrics & systems pharmacology, 6(8):499–511, 2017.

[90] Dénes Türei, Tamás Korcsmáros, and J. Saez-Rodriguez. Omnipath: guidelines and gateway for literature-curated signaling pathway resources. Nature Methods, 13:966–967, 2016.

[91] Charles J Vaske, Stephen C Benz, J Zachary Sanborn, Dent Earl, Christopher Szeto, Jingchun Zhu, David Haussler, and Joshua M Stuart. Inference of patient-specific pathway activities from multi-dimensional cancer genomics data using paradigm. Bioinformatics, 26(12):i237–i245, 2010.

[92] Ashish Vaswani, Noam Shazeer, Niki Parmar, Jakob Uszkoreit, Llion Jones, Aidan N. Gomez, Lukasz Kaiser, and Illia Polosukhin. Attention is all you need, 2023.

[93] Petar Velickovic, Guillem Cucurull, Arantxa Casanova, Adriana Romero, Pietro Lio’, and Yoshua Bengio. Graph attention networks. ArXiv, abs/1710.10903, 2017.

[94] Shangying Wang, Kai Fan, Nan Luo, Yangxiaolu Cao, Feilun Wu, Carolyn Zhang, Katherine A Heller, and Lingchong You. Massive computational acceleration by using neural networks to emulate mechanism-based biological models. Nature communications, 10(1):4354, 2019.

[95] James W. Wilson, Dilem Shakir, Michael Batie, Mark Frost, and Sonia Rocha. Oxygen-sensing mechanisms in cells. The FEBS Journal, 287(18):3888–3906, 2020.

[96] David S Wishart, Craig Knox, An Chi Guo, Savita Shrivastava, Murtaza Hassanali, Paul Stothard, Zhan Chang, and Jennifer Woolsey. Drugbank: a comprehensive resource for in silico drug discovery and exploration. Nucleic acids research, 34(suppl_1):D668–D672, 2006.

[97] David H. Wolpert and William G. Macready. No free lunch theorems for optimization. IEEE Trans. Evol. Comput., 1:67–82, 1997.

[98] Keyulu Xu, Weihua Hu, Jure Leskovec, and Stefanie Jegelka. How powerful are graph neural networks? ArXiv, abs/1810.00826, 2018.

[99] Keyulu Xu, Chengtao Li, Yonglong Tian, Tomohiro Sonobe, Ken ichi Kawarabayashi, and Stefanie Jegelka. Representation learning on graphs with jumping knowledge networks. ArXiv, abs/1806.03536, 2018.

[100] Qing-Song Xu and Yi-Zeng Liang. Monte carlo cross validation. Chemometrics and Intelligent Laboratory Systems, 56(1):1–11, 2001.

[101] Rex Ying, Dylan Bourgeois, Jiaxuan You, Marinka Zitnik, and Jure Leskovec. Gnnexplainer: Generating explanations for graph neural networks, 2019.

[102] Bo Yuan, Ciyue Shen, Augustin Luna, Anil Korkut, Debora S Marks, John Ingraham, and Chris Sander. Cell-box: interpretable machine learning for perturbation biology with application to the design of cancer combination therapy. Cell systems, 12(2):128–140, 2021.

[103] Chuxu Zhang, Dongjin Song, Chao Huang, Ananthram Swami, and Nitesh V. Chawla. Heterogeneous graph neural network. In Proceedings of the 25th ACM SIGKDD International Conference on Knowledge Discovery & Data Mining, KDD ‘19, page 793–803, New York, NY, USA, 2019. Association for Computing Machinery.

[104] Lingxiao Zhao and Leman Akoglu. Pairnorm: Tackling oversmoothing in gnns, 2020.

[105] Wei Zhao, Jun Li, Mei-Ju M Chen, Yikai Luo, Zhenlin Ju, Nicole K Nesser, Katie Johnson-Camacho, Christopher T Boniface, Yancey Lawrence, Nupur T Pande, et al. Large-scale characterization of drug responses of clinically relevant proteins in cancer cell lines. Cancer cell, 38(6):829–843, 2020.

[106] Xin Zheng, Yixin Liu, Shirui Pan, Miao Zhang, D. Jin, and Philip S. Yu. Graph neural networks for graphs with heterophily: A survey, 2022.

[107] Jiong Zhu, Yujun Yan, Lingxiao Zhao, Mark Heimann, Leman Akoglu, and Danai Koutra. Beyond homophily in graph neural networks: Current limitations and effective designs, 2020.

